# Small molecule targeting of FBXO21 mediated p85α ubiquitylation in acute myeloid leukemia

**DOI:** 10.1101/2024.12.13.628427

**Authors:** Kasidy K. Dobish, Suchita Vishwakarma, Hendrik C. Peters, C. Bea Winship, Danielle Alvarado, R. Katherine Hyde, Amarnath Natarajan, Shannon M. Buckley

## Abstract

PI3K inhibitors that target the catalytic sub-unit p110 are used in cancer therapy to inhibit overactive PI3K signaling pathway. However, their clinical use is limited by severe adverse effects and development of resistance, highlighting the need for further research on regulation of the PI3K pathway. We have identified that FBXO21 ubiquitinates regulatory PI3K subunit p85α, and silencing of FBXO21 inhibits canonical PI3K signaling. To target FBXO21, we developed a small molecule designed to interfere with substrate:ligase interaction. Our novel small molecule effectively blocks p85α ubiquitination leading to decreased PI3K pathway activation and cell death in acute myeloid leukemia (AML). Moreover, our studies demonstrate selectivity for AML cells over healthy counterparts, and elimination of AML *in vivo*, emphasizing FBXO21’s potential as a promising therapeutic target for AML. Targeting substrate:ligase interactions provide new avenues for drug discovery that may enhance the efficacy of current therapies and benefits for improving patient outcomes.

**STATEMENT OF SIGNIFICANCE:** Our studies highlight the potential of the ubiquitin E3 ligase FBXO21 as an alternative therapeutic target for the PI3K signaling pathway, not only in AML but in cancers with aberrant PI3K signaling.

## INTRODUCTION

The breakdown of regulatory proteins through the ubiquitin-proteasome system plays a crucial role in controlling the activation or deactivation of a cell’s molecular machinery at an appropriate time and within the correct subcellular compartment. The precision of this system is dependent on numerous ubiquitin ligases, with the SKP1-CUL1-FBOX (SCF) family of RING-finger E3 ligases being the largest group.^1,2^. Notably, approximately 70 distinct FBOX proteins have been identified.^3^ All FBOX proteins are categorized into three groups based on their substrate binding domain: FBXW (F-box with WD40 domains), FBXL (F-box with leucine-rich repeats), and FBXO (F-box proteins with other domains). One characteristic shared by most FBOX proteins is the need for a phosphodegron. A phosphodegron is a specific amino acid sequence, typically containing a serine or a threonine or a tyrosine, whose phosphorylation permits recognition by the E3 ligase preceding ubiquitination for proteasomal degradation. These sequences can be unique or have specific commonalities. An example includes the TPxxS degron motif for FBXW7 (LLPTPPLS is the c-MYC degron motif).^4,5^ FBOX proteins recognize about 20% of proteins degraded by the UPS and have been found to ubiquitinate substrates important for cell survival, cell cycle regulators, transcription factors, and cell-surface receptors.^6,7^ The FBOX substrate interface has recently been the focus of drug discovery efforts.^8^ For instance, substituted-aryl-ethane-1,2-diamine analogs (BC1215 and BC1258) inhibit FBXO3-Fbxl2 interaction to either stabilize TRAF or promote Aurora B degradation, indicating tractability of targeting FBOX proteins.^9,10^ These studies emphasize the critical role of FBOX E3 ligases as targetable proteins for drug discovery to create more effective and targeted therapies.

We found high expression levels of the FBOX ubiquitin E3 ligase, FBXO21, is associated with poor patient survival in acute myeloid leukemia (AML). Silencing *FBXO21* in AML leads to differentiation, and delayed tumor progression. Further, we identified p85α, a regulatory subunit of the phosphoinositide 3-kinase (PI3K) pathway as a novel substrate of FBXO21, marking it for proteasomal degradation. Depletion of FBXO21 stabilizes p85α, leading to dimerization of free p85 and decreased activation of the PI3K pathway.^11^ Therefore, FBXO21 may serve as a novel regulator of the PI3K signaling pathway, offering an alternative approach to modulating PI3K activity in AML. While clinical use of PI3K inhibitors for targeting the PI3K/AKT signaling pathway in cancer treatment is limited by off-target effects, dose-limiting toxicities, and resistance mechanisms, targeting FBXO21 may provide a viable alternative.^12^ Depleting *FBXO21* in healthy hematopoietic stem and progenitor cells (HSPCs) *in vitro* and in *Fbxo21* knockout mouse models showed minimal to no impact on normal hematopoiesis, suggesting that targeting FBXO21 may selectively target the PI3K system in malignant cells without the associated toxicity seen with traditional PI3K inhibitors.^13^

FBXO21 interacts with the previously identified substrate, EID1, through its YccV domain.^14^ YccV domains within proteins are characteristically highly conserved, indicating their importance in many cellular functions.^15^ By characterizing specifics of the FBXO21/p85α interaction, which requires the YccV domain of FBXO21 and the phosphorylation of the Y467 residue within the iSH2 domain of p85α, we gain crucial insights into how to effectively interfere with this interaction. This detailed understanding allowed for the development of a targeted therapy that can disrupt the FBXO21/p85α interaction. We developed a terphenyl analog as a helical mimic that interacts with the substrate recognition domain to block FBXO21 mediated p85α ubiquitination, resulting in decreased PI3K pathway activation in AML cells. Targeting the ubiquitin E3 ligase FBXO21 and its regulation of p85α ubiquitylation offers a novel approach to modulating the PI3K signaling pathway in AML while having little to no toxicities on healthy cells.

## RESULTS

### FBXO21 mediates K-48 linked polyubiquitination of p85α at K692

To determine the specificity of p85α ubiquitination by FBXO21, a panel of F-box proteins was screened using a cell-free *in vitro* ubiquitination assay. Our screen revealed that only FBXO21 was able to facilitate the ubiquitination of p85α (Figure 1A). Consistent with previous studies showing FBXO21 interacts with previously identified substrate, EID1, through its YccV domain, we found that ubiquitination of p85α depends on the YccV domain (residues 503-557) of FBXO21 (Figure 1B).^14^ *In vitro* ubiquitination assay demonstrated immunopurified FBXO21, but not the YccV-deficient FBXO21 mutant, was able to ubiquitinate p85α *in vitro* (Figure 1B). This suggests that the YccV domain of FBXO21 serves as the substrate recognition domain required for binding to p85α.

**Figure 1:**
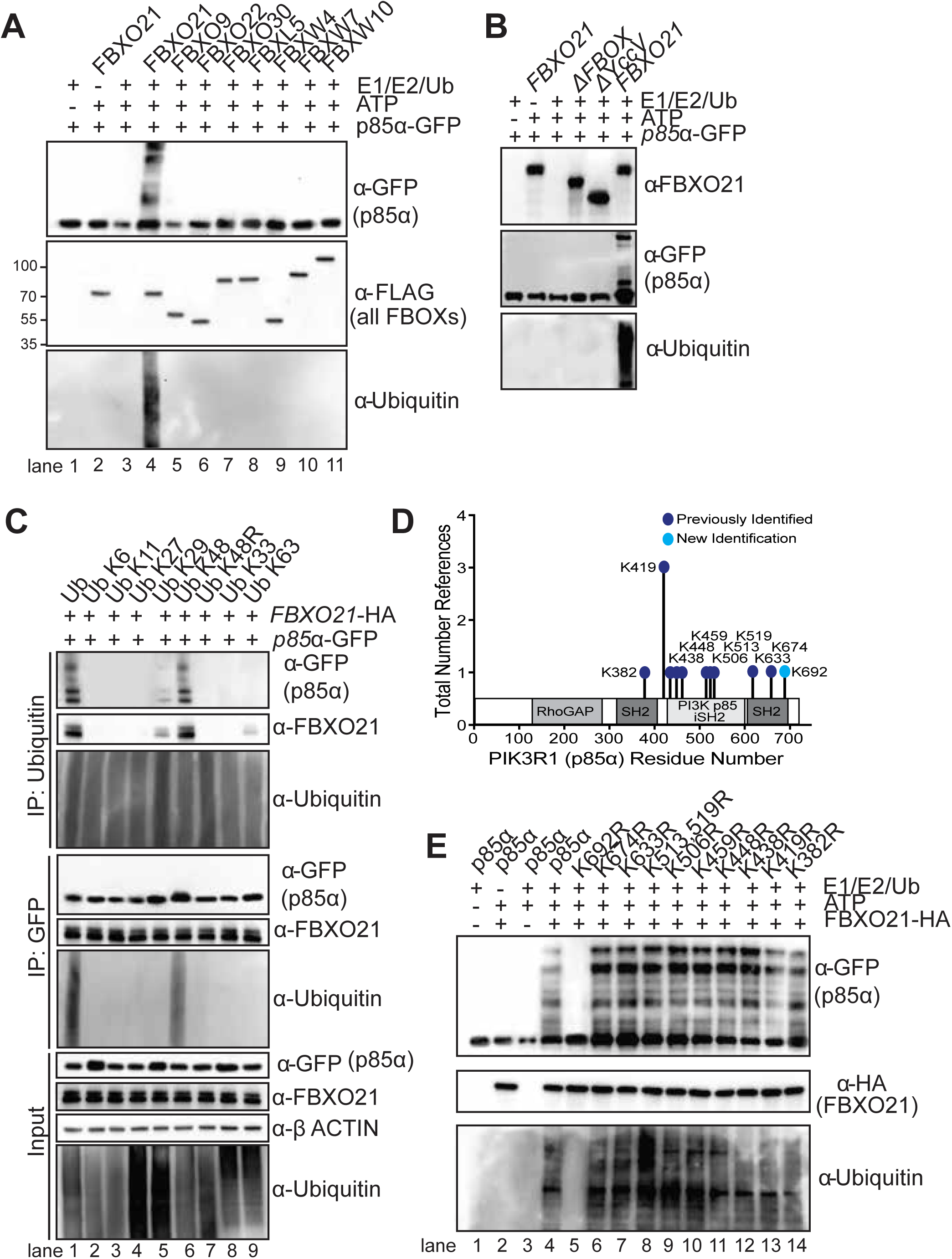
FBXO21 specifically ubiquitinates p85α through K48-linked ubiquitination, requiring the YccV domain and lysine residue K692. (n=3 for all experiments) **A** Western blot of *in vitro* ubiquitination assay of GFP-tagged *p85α* and FLAG-tagged FBXO21, FBXO9, FBXO22, FBXO30, FBXL5, FBXW4, FBXW7, or FBXW10 in the presence of E1, E2, E3 (FLAG-FBOX protein) and ubiquitin (Ub). **B** Western blot of *in vitro* ubiquitination assay including GFP-tagged *p85α*, HA-tagged *FBXO21*, HA-tagged *ΔFBXO21*, and HA-tagged *FBXO21-ΔYccV* was performed in the presence of E1, E2, and ubiquitin (Ub). **C** Western blot of Ubiquitin and GFP (*p85α*) immunoprecipitation in HEK293T cells transiently transfected with HA-tagged *FBXO21,* GFP-tagged *p85α,* and wild-type Ubiquitin or Ubiquitin lysine mutant K6, K11, K27, K29, K48, K48R, K33, or K63. **D** Schematic depicting the previously identified sites of ubiquitination and new lysine residue identified by K-ε-GG immunoprecipitation followed by mass spectrometry. **E** Western blot of *in vitro* ubiquitination assay of GFP-tagged WT *p85α* and GFP-tagged *p85α* with lysine to arginine mutation at each identified ubiquitination site in the presence of E1, E2, E3 (HA-FBXO21) and ubiquitin (Ub).

To further explore the characteristics of this ubiquitination and the lysine linkage required for FBXO21 ubiquitination of p85α, we transiently expressed HA tagged wild-type (WT) ubiquitin or a ubiquitin lysine (K) mutant. The K linkage mutants only contained the lysine linkage specified, with all other lysines mutated to arginine (R). The ubiquitin WT or mutants were transfected in addition to FLAG-tagged FBXO21 and GFP-tagged p85α. We immunoprecipitated GFP or HA and found that ubiquitinated p85α protein was only detected in cells expressing WT ubiquitin and K48-linked ubiquitin, but not in cells with a K48 to arginine (R) mutation (Figure 1C). This suggests that FBXO21 specifically ubiquitinates p85α through K48-linked ubiquitination. Ten lysine residues have previously been identified as sites of ubiquitination on p85α.^16^ We identified a new lysine residue, K692, from our K-ε-GG immunoprecipitation followed by mass spectrometry (Figure 1D)^11,17–19^ as the target of FBXO21 mediated p85α ubiquitination. Immunopurified p85α-K692R (lane 5) led to the loss of ubiquitination, while the other ten previously reported lysine mutations did not affect p85α ubiquitination, demonstrating that FBXO21 mediates polyubiquitination of p85α selectively at K692 (Figure 1E). These findings provide important insights into the molecular mechanisms underlying the ubiquitination of p85α by FBXO21 and further our understanding of the regulatory role of FBXO21 in the modulation of the PI3K signaling pathway.

### p85α phosphodegron within iSH2 domain required for FBXO21 binding

Next, we determined the FBXO21-binding motif in p85α by using deletion mutants to identify the domain of p85α required for binding (Figure 2A). We transiently transfected GFP-tagged domain mutants of p85α alongside HA-tagged FBXO21 into HEK293T cells and performed immunoprecipitation for both FBXO21 and p85α. Interestingly, we found that the interaction between p85α and FBXO21 was lost upon deletion of the iSH2 domain of p85α (Figure 2B). The iSH2 (inter-SH2) domain of p85α directly binds to the p110 catalytic subunit of PI3K functioning as a tether to maintain the structural integrity of the p85α/p110 complex, which is essential for the activation and proper functioning of the PI3K pathway.^20^ To identify the sequence in p85α that interacts with FBXO21, we conducted comparative bioinformatics analyses to predict small peptide sequences that had a high probability of interacting with the YccV domain of FBXO21. Three potential degron sequences, L^466^YEEYT^471^, I^493^FEEQC^498^, and R^544^LEEDL^549^ on p85α were identified (Figure 3A). The top three identified sites were all within the iSH2 domain of p85α, further confirming the results of our immunoprecipitation. Overlay of the identified docked p85α sequences with EID1 sequence (F^161^IEELF^166^) identified L^466^YEEYT^471^ in p85α showed comparable docking scores (Figure 3B). It is typical for FBOX ubiquitin E3 ligases to require phosphorylation of their substrates, or phosphorylation of an amino acid within their specific degron, for recognition. Only one of three identified predicted binding sequences (L^466^YEEYT^471^) included an amino acid that can undergo phosphorylation (i.e. tyrosine (Y), threonine (T), and serine (S)).

**Figure 2:**
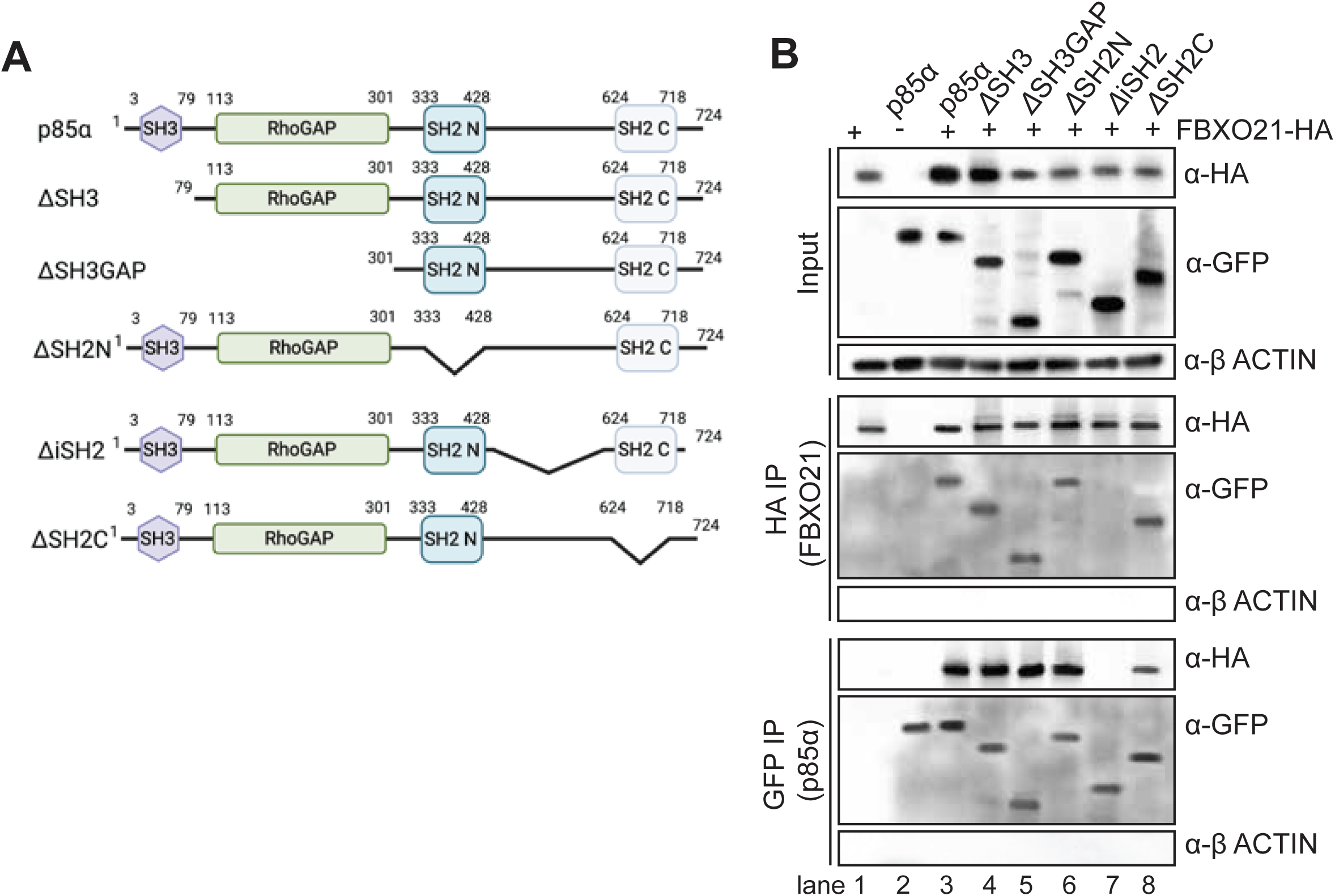
FBXO21 binds to and ubiquitinates p85α via the iSH2 domain and phosphorylation site Y467. (n=3 for all experiments) **A** Schematic representation of p85*α* mutants **B** Western blot of HA (FBXO21) and GFP (*p85α*) immunoprecipitation in HEK293T cells transiently transfected with HA-tagged *FBXO21* and WT or mutated GFP-tagged *p85α*

**Figure 3:**
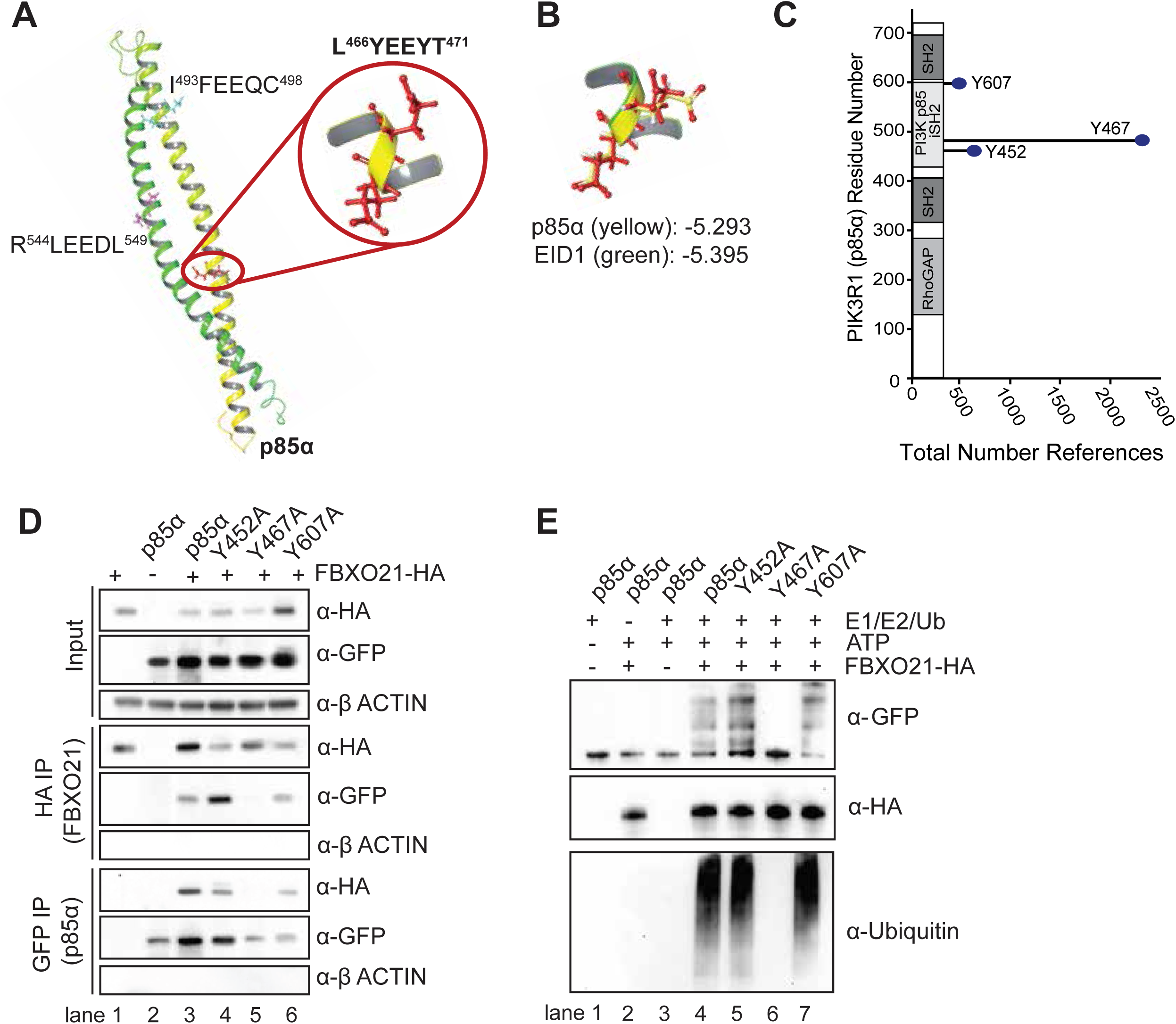
Phosphodegron containing Y467 is required for p85α ubiquitination. (n=3 for all experiments) **A** The secondary structure of p85α is shown as ribbon representation. Comparative bioinformatics analyses identified the highlighted sequences on p85α that had a high probability of interacting with FBXO21 YccV domain. **B** Overlay of the docked (Schrodinger’s Glide) p85α sequences predicted L^466^YEEYT^471^ as the FBXO21 binding p85α sequence. **C** Schematic depicting selected previously identified sites of phosphorylation on p85*α.* **D** Western blot of HA (FBXO21) and GFP (*p85α*) immunoprecipitation in HEK293T cells transiently transfected with HA-tagged *FBXO21* and WT or tyrosine (Y) to alanine (A) mutated GFP-tagged *p85α.* **E** Western blot of *in vitro* ubiquitination assay of GFP-tagged WT *p85α* and GFP-tagged *p85α* with tyrosine (Y) to alanine (A) mutation at each identified phosphorylation site in the presence of E1, E2, E3 (HA-FBXO21) and ubiquitin (Ub).

Utilizing the predicted computational modeling and previously identified phosphorylation sites, we mutated top phosphorylated tyrosine (Y) sites within the iSH2 domain of p85α to alanine (A), rendering these residues incapable of phosphorylation (Figure 3C).^16^ We found that FBXO21 mediated p85α ubiquitination was lost only in the Y467A p85α mutant, suggesting that phosphorylation of Y467 is required for FBXO21 mediated degradation of p85α (Figure 3D-E). Together, we have defined the molecular basis for FBXO21 mediated p85α ubiquitination, which indicates that a helical mimic of p85α LYEEYT would bind to the YccV domain of FBXO21 to disrupt p85α ubiquitination.

### Small molecule inhibits p85α ubiquitination

The region of p85α that binds to the YccV domain of FBXO21 adopts an α-helical conformation (Figure 4A). While there are several approaches to design α-helical mimics, we focused on non-peptide α-helix mimetics.^21–26^ Specifically, we were inspired by the elegant use of substituted terphenyl analogs as α-helix mimetics that was pioneered by the Hamilton group.^27–29^ We initiated an iterative design, dock, synthesize and evaluate (DDSE) approach to identify a substituted terphenyl analog that would disrupt FBXO21 mediated ubiquitination of p85α. The binding mode of the active compound against FBXO21 (57-057) is supported by structure activity relationship of related analogs (Figure 4B). The FBXO21 mediated *in vitro* ubiquitination of p85α was inhibited by the terphenyl analogs 57-057 and 57-112. The model indicated that the carboxylic acid group on the active compounds was within hydrogen bond distance of R^561^ and H^511^. The complete loss of activity with 57-139 in which the carboxylic acid is replaced with a methyl group and 57-048 in which the carboxylic acid was converted to the ester, suggests that the interaction between the carboxylic acid in the active compounds and R^561^ and H^511^ in FBXO21 is critical. The model also indicates that the fluoro-phenyl ortho to the carboxylic acid occupies a narrow binding pocket, consistently replacing the fluorine atom with two atom nitrile group in 57-125 resulted in complete loss of activity.

**Figure 4:**
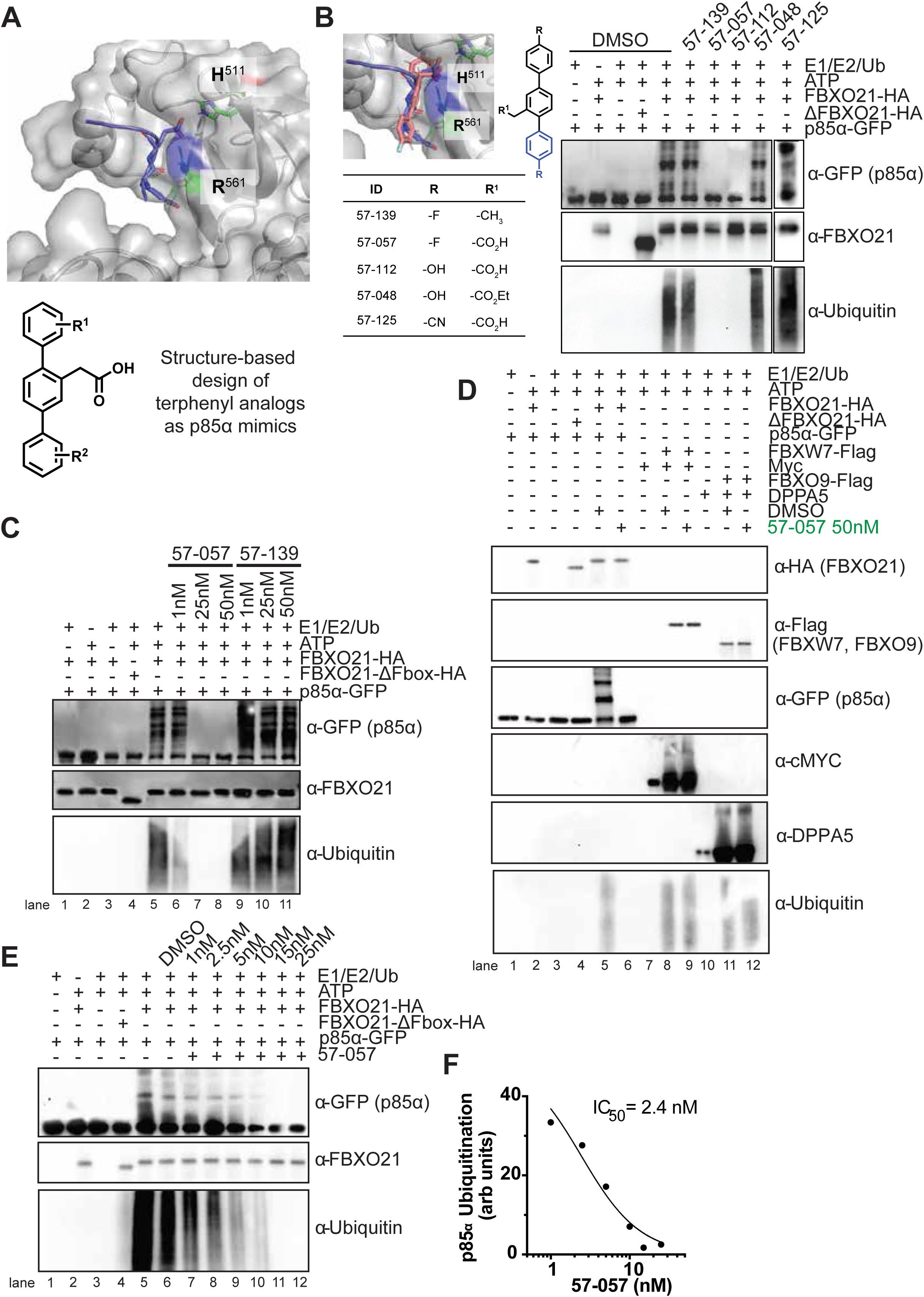
Inhibition of FBXO21/p85α interaction by terphenyl analog 57-057. (n=2 for all experiments) **A** Docked structure of p85α peptide (L<u>YE</u>EYT – <u>blue stick</u>) into YccV domain of FBXO21. Generic structure of terphenyl analogs used to identify p85a mimics. **B** Binding mode of FBXO21 inhibitor overlayed with the docked p85α peptides and the associated structure activity relationship that supports the model. **C** Western blot of an *in vitro* ubiquitination assay with 57-057 compound and its respective inactive compound 57-139 with concentrations ranging from 1 nM to 50 nM in the presence of E1, E2, and ubiquitin. **D** Western blot of *in vitro* ubiquitination showing the effects of 57-057 on the ubiquitination of other FBOX substrates. The substrates analyzed include cMYC for FBXW7 and DPPA5 for FBXO9, in the presence of their respective FBOX proteins and E1, E2, ATP, and ubiquitin. **E** Dose-response analysis of 57-057 in an i*n vitro* ubiquitination assay with concentrations ranging from 1 nM to 25 nM. The assay shows that 57-057 effectively inhibits the ubiquitination of p85α at low nanomolar concentrations. **F** Dose-response curve of 57-057 in inhibiting p85α ubiquitination with a cell-free IC50 value of 2.4 nM.

Follow up dose response studies with the active compound 57-057 inhibited FBXO21 mediated ubiquitination of p85α at low nanomolar concentrations (lanes 6-8), whereas the control compound 57-139 (lanes 9-11) did not (Figure 4C). This confirms that 57-057 effectively disrupts the FBXO21/p85α interaction. Next, we assessed the selectivity of 57-057 in three *in vitro* ubiquitination assay systems *viz.*, FBXO21 mediated p85α ubiquitination, FBXW7 mediated cMyc ubiquitination,^30^ and FBXO9 mediated DPPA5 ubiquitination.^31^ Remarkably, we observed selective inhibition of FBXO21 mediated p85α ubiquitination by 57-057 (lane 5-6, Figure 4D). No such inhibition was observed in either FBXW7 mediated cMyc ubiquitination (lane 8-9, Figure 4D) or FBXO9 mediated ubiquitination of DPPA5 (lane 11-12, Figure 4D). Together, the data shows that 57-057 is a selective FBXO21 binder that inhibits the *in vitro* ubiquitination of p85α at low-nM potencies with an IC_50_ of 2.4 nM (Figure 4E-F).

The *in vitro* results were validated in AML models with p85α levels as the read out for small molecule mediated perturbation of FBXO21. Disruption of p85α binding to FBXO21 should prevent SCF mediated ubiquitination of p85α thus resulting in the stabilization / increase of p85α levels in cells. Consistent with our previous FBXO21 knockdown studies that resulted in increased p85α in AML cells,^32^ we observed a dose-dependent increase in p85α levels in MOLM-13 cells treated with 57-057. This disruption led to a decrease in AKT activation and a reduction in p85/p110 dimerization, accompanied by an increase in p85/p85 dimerization, consistent with the results observed from FBXO21 knockdown experiments (Supplemental Figure 1A-C).

### 57-057 shows selectivity to AML tumor cells over healthy HSPCs

Next, we tested the antiproliferative effects of 57-057 on AML cells, specifically MOLM-13 and HL-60. The compound induced a decrease in AML cell viability at low nanomolar concentrations, with IC_50_ values of 25 nM for MOLM-13 and 32 nM for HL-60 (Supplemental Figure 1A). A common challenge with PI3K inhibitors is the development of resistance. To test the efficacy of our compound against a PI3K inhibitor-resistant cell line, we created MOLM-13 Alpelisib-resistant cell lines. Alpelisib works by inhibiting the PI3K catalytic subunit, p110α, which is often mutated and hyperactivated in various cancers.^33,34^ Alpelisib specifically binds at the ATP-binding pocket of p110α, preventing the phosphorylation of PIP2 to PIP3, a crucial step in the PI3K/AKT signaling pathway.^35^ By blocking this enzyme, Alpelisib reduces the downstream signaling through the AKT/mTOR pathway, thereby inhibiting cancer cell growth and survival.^35^ Although PI3K inhibitors are not FDA approved for AML, the PI3K inhibitor Alpelisib is FDA approved for the treatment of HR-positive, HER2-negative advanced or metastatic breast cancer.^33,34^ We created MOLM-13 resistant cell lines by gradually exposing the cells to increasing concentrations of Alpelisib over several weeks until resistance was established, and confirmed by an increase in IC_50_ value from 115nM to 455nM (Supplemental Figure 1D). The increase in IC_50_ suggests that MOLM-13 cells have developed resistance to Alpelisib, which could be due to the activation of alternative survival pathways, such as MAPK or mTOR, which can bypass PI3K inhibition.^36,37^

We then tested 57-057 in both normal and Alpelisib-resistant MOLM-13 cells (Supplemental Figure 1E). Remarkably, even with resistance to the PI3K inhibitor targeting the p110α catalytic subunit, the small molecule targeting FBXO21 (57-057) remained effective (IC_50_ = 34nM). The accumulation of p85α and decreased canonical PI3K signaling in both normal and Alpelisib-resistant AML cells demonstrate that targeting FBXO21 leads to cell death through the same mechanism (Supplemental Figure 1F). This response suggests that targeting FBXO21 can circumvent the resistance mechanisms that often develop against traditional PI3K inhibitors. These findings demonstrate the potent and selective action of 57-057, as it inhibited the growth of AML cells by disrupting the FBXO21-p85α interaction and downregulating canonical PI3K signaling. The consistent efficacy in both normal and PI3K inhibitor-resistant cells highlights the therapeutic potential of targeting FBXO21 as a novel strategy in AML treatment.

Previously, we demonstrated that FBXO21 is dispensable for steady-state hematopoiesis but plays a critical role in the proliferation and survival of AML through its ubiquitination of p85α.^32,38^ Building on these results, we extended our investigation to primary healthy CD34+ HSPC and AML patient cells to evaluate the broader applicability and therapeutic potential of the 57-057 compound. We found 57-057 exhibited approximately 5-fold selectivity (8 nM vs. 39 nM) for killing primary AML cells over CD34+ HSPCs (Figure 5A). In contrast, Alpelisib did not show such selectivity (47 nM vs. 56 nM) (Figure 5B). In direct comparison with Alpelisib we found 57-057 inhibited the growth of primary AML cells with approximately 6-fold higher potency (8 nM vs. 47 nM) (Figure 5C). We also observed increased p85α accumulation in both AML and CD34+ cells treated with the active compound (57-057), validating that our compound is blocking the ubiquitination of p85α by FBXO21 (Figure 5D). There was also a corresponding decrease in AKT activation, which was more apparent in AML cells in comparison to healthy CD34+ cells (Figure 5D). The greater decrease in AKT activation in AML cells compared to healthy cells upon 57-057 treatment reflects both the higher dependency of cancer cells on the PI3K/AKT pathway for their growth and survival, and the selective cytotoxicity of 57-057 which targets the hyperactive PI3K/AKT pathway by blocking the ubiquitination of p85α, leading to cytotoxic effects predominantly in cancer cells. Interestingly, we noted a dose-dependent increase in p85α /p85α dimerization in both CD34+ and AML samples, and a corresponding loss of p85α /p110 dimerization in AML samples (Figure 5E). Furthermore, in colony-forming assays with primary AML and healthy CD34+ cells treated with increasing concentrations of 57-057, we found no significant decrease in colony formation by CD34+ cells treated with 57-057. Importantly, 57-057 treatment in primary AML cells resulted in a dramatic dose-dependent decrease in the number of colonies formed, suggesting a loss of stemness and reduced growth rate (Figure 5F). These findings underscore the potential of 57-057 as a more effective and selective therapeutic option for targeting AML cells, including those resistant to conventional PI3K inhibitors.

**Figure 5:**
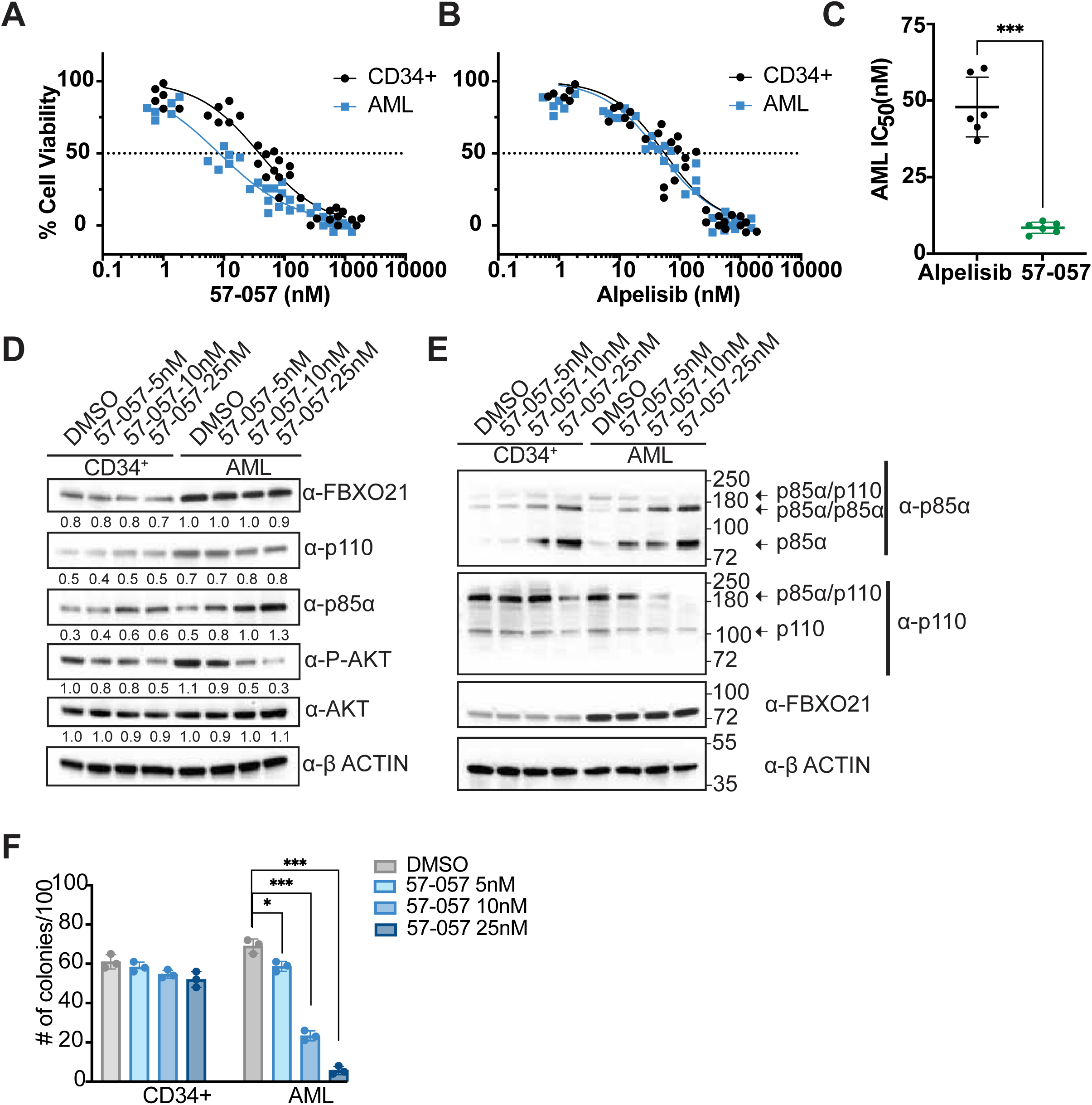
Efficacy and specificity of 57-057 in primary AML and healthy CD34^+^ cells. (n=3 independent CD34^+^ patient samples, n=4 independent AML patients) MTT assays were performed for 72 hours to assess the viability of primary AML cells and healthy CD34+ cells treated with varying concentrations of **A** 57-057 and **B** Alpelisib. **C** IC50 values for 57-057 and Alpelisib in primary AML cells, highlighting the significantly lower IC50 for 57-057. **D** Western blot analysis of FBXO21, p110, p85α, P-AKT, AKT, and β-ACTIN and **E** native gel analysis for PI3K complex proteins in primary AML cells and CD34+ cells treated with varying concentrations between 5-25nM of 57-057 or DMSO for 72 hours. **F** Colony-forming assays of primary AML and CD34+ cells treated with increasing concentrations of 57-057 between 5-25nM, or equal volume of DMSO control. (** p ≤ 0.01, *** p ≤ 0.001).

### 57-057 shows promise for AML treatment through effective targeting of FBXO21 and favorable pharmacokinetics

To advance the compound 57-057 for further study, we first tested it in MLL-AF9 mouse spleen tumors to determine if it would be as effective on mouse AML cells as it is on human AML cells.^39^ We confirmed that treating these tumor cells for 72 hours *in vitro* with increasing doses of 57-057 resulted in increased apoptosis, decreased colonies, and decreased canonical PI3K activity, as evidenced by reduced AKT phosphorylation (Figure 6A-C). Pharmacokinetic (PK) and tissues distribution studies reveal that 57-057 (30 mg/kg) is orally bioavailable, with a half-life of ∼6h (Supplemental Figure 2A) and is primarily cleared by the liver (Supplemental Figure 2B).

**Figure 6:**
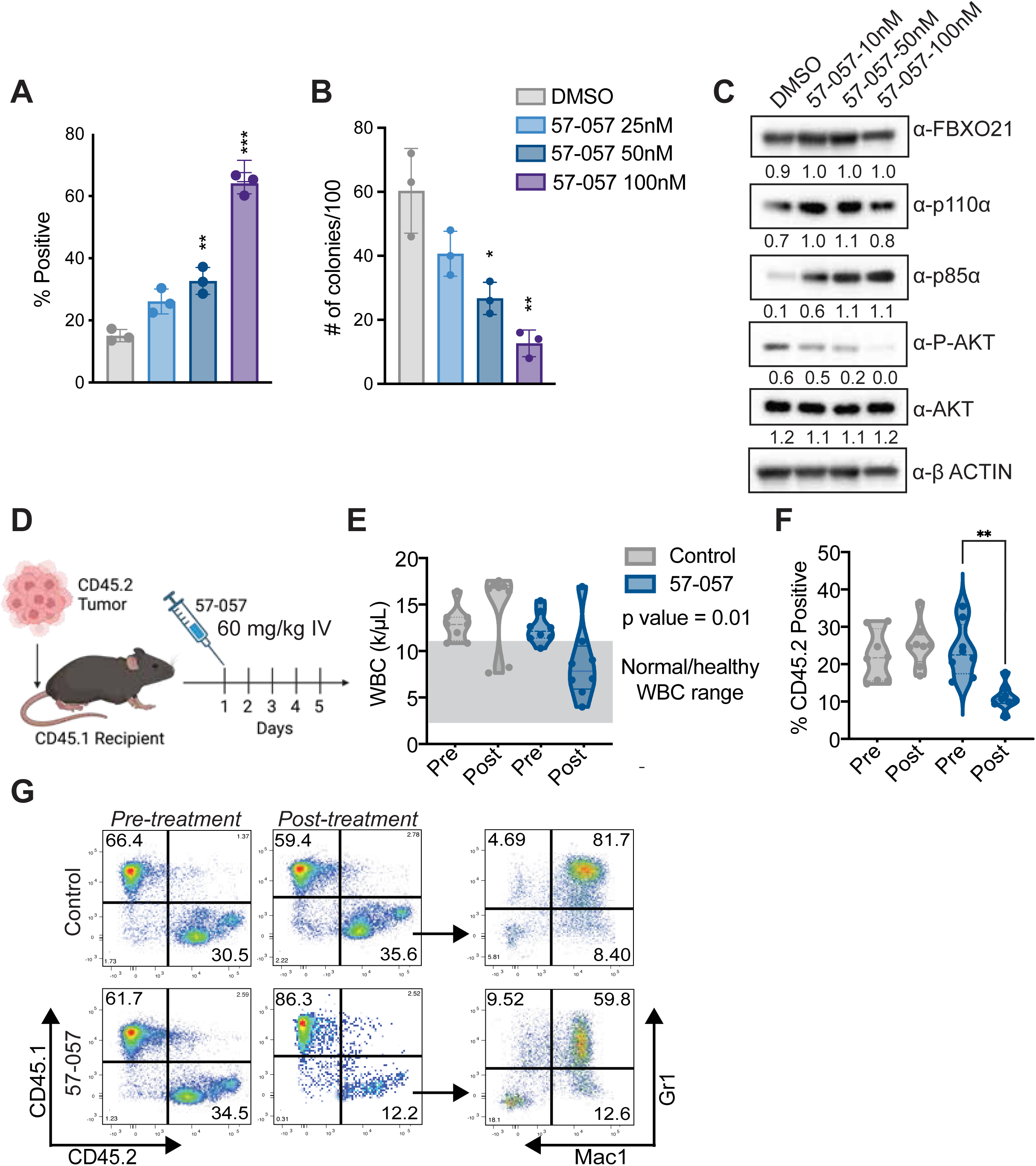
57-057 targets AML tumors in mouse model of MLL-AF9. MLL-AF9 spleen tumor cells were cultured and treated with a novel FBXO21 inhibitor, compound 57-057, for 72 hours and analyzed for **A** apoptosis by flow cytometry, **B** colony formation, and **C** Western blot showing the effects of 57-057 on the expression levels of FBXO21, p110α, p85α, P-AKT, AKT, and β-ACTIN (n=2). **D** Schematic representation of the experimental design. CD45.2 MLL-AF9 AML spleen tumor cells were transplanted into sub lethally irradiated CD45.1 recipient mice. Tumor engraftment was confirmed, and mice were treated with 57-057 at 60 mg/kg by IV injection for five consecutive days. Mice were sacrificed 72 hours post-treatment for analysis. **E** White blood cell (WBC) counts and **F** flow cytometric analysis of CD45.2 positive cells pre- and post-treatment in control and 57-057 treated mice. **G** Representative flowcytometry plot pre- and post-treatment.

Dose escalation studies treating mice IP starting at 2.5mg/kg to 90mg/kg over a 14-week period showed no adverse effects with 57-057 treatment (normal white blood cell counts, no weight loss, and altered metabolic enzymes). Further, we observed no alterations in the percentage of long-term hematopoietic stem cells (LT-HSCs), short-term hematopoietic stem cells (ST-HSCs), or any multipotent progenitors (MPPs), common myeloid progenitors (CMPs), granulocyte-monocyte progenitors (GMPs), or megakaryocyte-erythroid progenitors (MEPs), indicating that 57-057 does not impair stem or progenitor populations *in vivo* (Supplemental Figure 2C). Additionally, there were no differences in bone marrow count or spleen weight across the treatment groups. (Supplemental Figure 2D-E). This is crucial as it suggests that 57-057 does not indiscriminately target healthy cells, preserving normal hematopoiesis and reducing the risk of bone marrow suppression.

Next, to evaluate the efficacy of 57-057 in an AML disease model, we conducted a transplantation study where CD45.2 MLL-AF9 AML spleen tumor cells were transplanted into sub-lethally irradiated CD45.1 recipient mice. After confirming tumor engraftment and progression of leukemia, we treated half of these mice with 60 mg/kg of 57-057 by IV injection for five consecutive days and sacrificed for analysis 72 hours post-final treatment. Overall, the treated group showed decreased WBC count and reduced leukemic burden, as evidenced by a decrease in tumor cell engraftment and a reduction in the Gr1/Mac1 positive tumor population, which represents myeloid leukemia cells critical for disease progression (Figure 6D-G). Collectively, our data demonstrates that 57-057 effectively reduces both the overall leukemic burden and specific leukemic cell populations, highlighting its potential as a targeted therapy that can selectively impact AML cells while minimizing harm to normal cells.

## DISCUSSION

The findings elucidate the critical role of FBXO21 in the ubiquitination and subsequent proteasomal degradation of p85α, providing new insights into the regulation of the PI3K signaling pathway in AML. Our previous studies demonstrated that FBXO21 minimally impacts steady-state hematopoiesis or the maintenance of hematopoietic stem cells, yet its overexpression in AML correlates with poor prognoses.^11,13^ This mechanism is mediated through the ubiquitination of p85α by FBXO21, leading to downstream alterations in PI3K signaling. Among various FBOX proteins, only FBXO21 could facilitate the ubiquitination of p85α, emphasizing its unique role and specificity in targeting p85α. Further, we identified lysine site K692 on p85α as a novel ubiquitination site using a K-ε-GG immunoprecipitation followed by mass spectrometry. Mutation of lysine residue K692R on p85α resulted in the loss of ubiquitination, confirming that FBXO21 specifically mediates ubiquitination at this newly identified site, providing a precise molecular target for potential therapeutic interventions aimed at modulating p85α stability. This highlights the utility of advanced proteomic approaches like K-ε-GG IP and mass spectrometry in uncovering critical post-translational modification sites that can be leveraged for targeted cancer therapies.

We found the YccV domain of FBXO21 to be the essential substrate binding domain, which is consistent with reported studies. Deletion of this domain abrogated the ability of FBXO21 to ubiquitinate p85α. Computational modeling identified key degron sequence L^466^YEEYT^471^ within p85α’s iSH2 domain necessary for FBXO21 binding. Loss of activity as a result of mutation of the phosphorylation site (Y467A) suggests that p85α could be phosphorylated at this site and FBXO21 binding followed by p85α ubiquitination is phosphorylation dependent. Understanding these precise molecular mechanisms is critical for the development of selective inhibitors that can modulate protein stability and function in cancer cells. The identification of other phospho-degrons has significantly advanced drug discovery efforts. For instance, elucidating the phosphodegron of IκBα for ubiquitination by β-TrCP has been pivotal in developing inhibitors that modulate NF-κB signaling, and the phosphodegron of IκBα served as the E3-ligase ligand for the first PROTAC.^40–42^

A structure-guided design led to the identification of a terphenyl analog, 57-057, which mimics the helical region of p85α. We show that 57-057 effectively disrupted FBXO21 mediated ubiquitination of p85α at low nM concentrations. This compound exhibited high selectivity for FBXO21 but not other FBOX proteins. In AML cell models 57-057 treatment resulted in stabilization of p85α, reduced AKT activation, and selectively inhibited the growth of cancer cells without affecting healthy cells. Our studies also suggest that targeting FBXO21 could overcome resistance mechanisms commonly encountered with PI3K inhibitors. In primary AML and CD34+ HSPC, 57-057 displayed selective growth inhibitory effects towards AML cells, with approximately 5-fold selectivity over healthy cells, and was ∼6-fold more potent than p110α inhibitor Alpelisib.

Pharmacokinetic studies revealed that 57-057 (30 mg/kg) is orally bioavailable, with a half-life of approximately 6 hours, primarily cleared by the liver. This half-life, while shorter compared to some therapies like Venetoclax, offers advantages in managing the drug’s safety profile by potentially reducing prolonged side effects and toxicity, such as cardiotoxicity. Remarkably, just 5 days of 57-057 treatment depleted over 60% of the tumor in the peripheral blood suggesting further *in vivo* efficacy studies, finetuning of dosing, and combination treatment could eradicate the tumor *in vivo*. In conclusion, our study highlights the critical role of FBXO21 in the regulation of p85α and the PI3K signaling pathway within hematopoietic malignancies and validates FBXO21 as a therapeutic target. The development of the selective compound against FBXO21, 57-057, offers a promising therapeutic strategy for targeting hematopoietic malignancies.

## MATERIALS AND METHODS

### *In Vitro* Ubiquitination

HEK293T cells were transfected with plasmids encoding HA-*FBXO21*, HA-*ΔFBXO21*, or GFP-*p85α,* with or without lysine(K) to arginine(R), or tyrosine(Y) to alanine(A) mutations. *FBXO21* and *ΔFBXO21* were a kind gift from Dr. Yukiko Yoshida, Tokyo Metropolitan Institute of Medical Science.^14^ 48hrs post transfection, HA-*FBXO21*, HA-*ΔFBXO21*, or GFP-*p85α* were immunopurified from the whole cell extracts using Anti-HA (Sigma) or Anti-GFP (MBL International) beads overnight at 4°C. The immunopurified HA-*FBXO21* or HA-*ΔFBXO21* (0.5 µg) proteins were incubated with immunopurified GFP-*p85α* (0.5 µg), E1 (500 ng), E2-UbcH5a (500 ng), FLAG-ubiquitin (0.5 µg) (BostonBiochem), and ATP (10mM). Ubiquitylation reactions were performed in 100 mM NaCl, 1 mM DTT, 5 mM MgCl2, 25 mM Tris-Cl (pH 7.5), incubated at 30°C for 2hrs, and stopped with 2x laemmli buffer (10 min at 95°C).

### Chemistry

Chemical synthesis of the compounds was performed as indicated in the Supplementary information. The synthesized compounds were characterized by ^1^HNMR, ^13^CNMR, ^19^FNMR, HRMS, GCMS and HPLC.

### Cell Culture

HEK293T, HL-60, and MOLM-13 cells were purchased from ATCC and DSMZ, and cultured according to ATCC/DSMZ guidelines. Primary human AML cells (Cureline Translational Cro) and CD34^+^ cells were cultured in StemSpan SFEM II media with CD34+ Expansion Supplement (StemCell Technologies) and UM729 (1 μM). Primary (CD34+ and AML) cells, along with the human-derived cell lines, were treated with varying doses of the compound 57-057, ranging from 5 nM to 100 nM, for a duration of 72 hours.

Mouse MLL-AF9 Lin-, Sca1+, cKit+ (LSK) cells were isolated from bone marrow from mice with AML (high WBC and Gr1/Mac1+) and cultured in media consisting of Opti-MEM Media, 10% FBS, 1% Pen/Strep, 50µM β-Mercaptoethanol, 50ng/mL Stem Cell Factor, 50ng/mL FLT3 Ligand (FLT3-L), 10ng/mL Interleukin-6 (IL-6), and 10ng/mL Interleukin-3 (IL-3). Antibodies listed in supplemental materials and methods.

### AML mouse model

For spleen tumor transplants, one day prior to irradiation, antibiotics were added to the water. Mice were sub-lethally irradiated with 500 cGy 4 – 12 hours prior to transplantation. Samples were prepared under sterile conditions and suspended in sterile PBS for injection. Tumor cells were transplanted via retro-orbital injection. The drug, 57-057, was resuspended in 5% DMSO and corn oil for oral gavage in dose escalation study and in a 5% DMSO, 35% PEG3000, and 60% water solution for intravenous injection. All pharmacokinetic (PK) testing was conducted using a formulation of 10% DMSO, 2% NMP, 50% PEG400, and 38% water.

### Statistical analysis

All experiments were performed in triplicate unless noted and statistical analyses were performed using unpaired two-tailed Student’s t-test assuming samples of equal variance. Error bars depict the standard deviation ±mean.

## AUTHOR CONTRIBUTIONS

K.K.D., S.V., A.N. and S.M.B. conceived and designed the experiments. K.K.D., S.V., H.P., C.B.W., and D.A performed experiments and analysis. K.K.D., S.V., A.N. and S.M.B. wrote the manuscript. All authors reviewed the manuscript before submission.

## ACKNOWLEDGEMENTS

This publication was supported by the Huntsman Cancer Institute at the University of Utah, supported by the National Cancer Institute of the National Institutes of Health (NIH) under award number P30CA042014.. S.M.B. is supported by the National Institutes of Health R37CA262635, and R01AI53090. A.N. is supported by the National Institutes of Health R01CA197999, R01CA260749, R01CA262664 and R01CA276846. R.K.H was supported by R01 CA244900. The UNMC core facilities are in part supported by P20GM121316 and P30CA036727. This research was supported by the State of Nebraska through the Pediatric Cancer Research Group part of the Child Health Research Institute.

## CONFLICTS OF INTEREST

The authors have no conflicts of interest related to this work.

## DATA AVAILABILITY

The remaining data needed to evaluate all conclusions are available within the Article and/or Supplementary Information.

## Notes

### Competing Interest Statement

The authors have declared no competing interest.

