## Supplemental Material for "Small molecule targeting of FBXO21 mediated p85α ubiquitylation in acute myeloid leukemia"

### Supplementary Figure 1

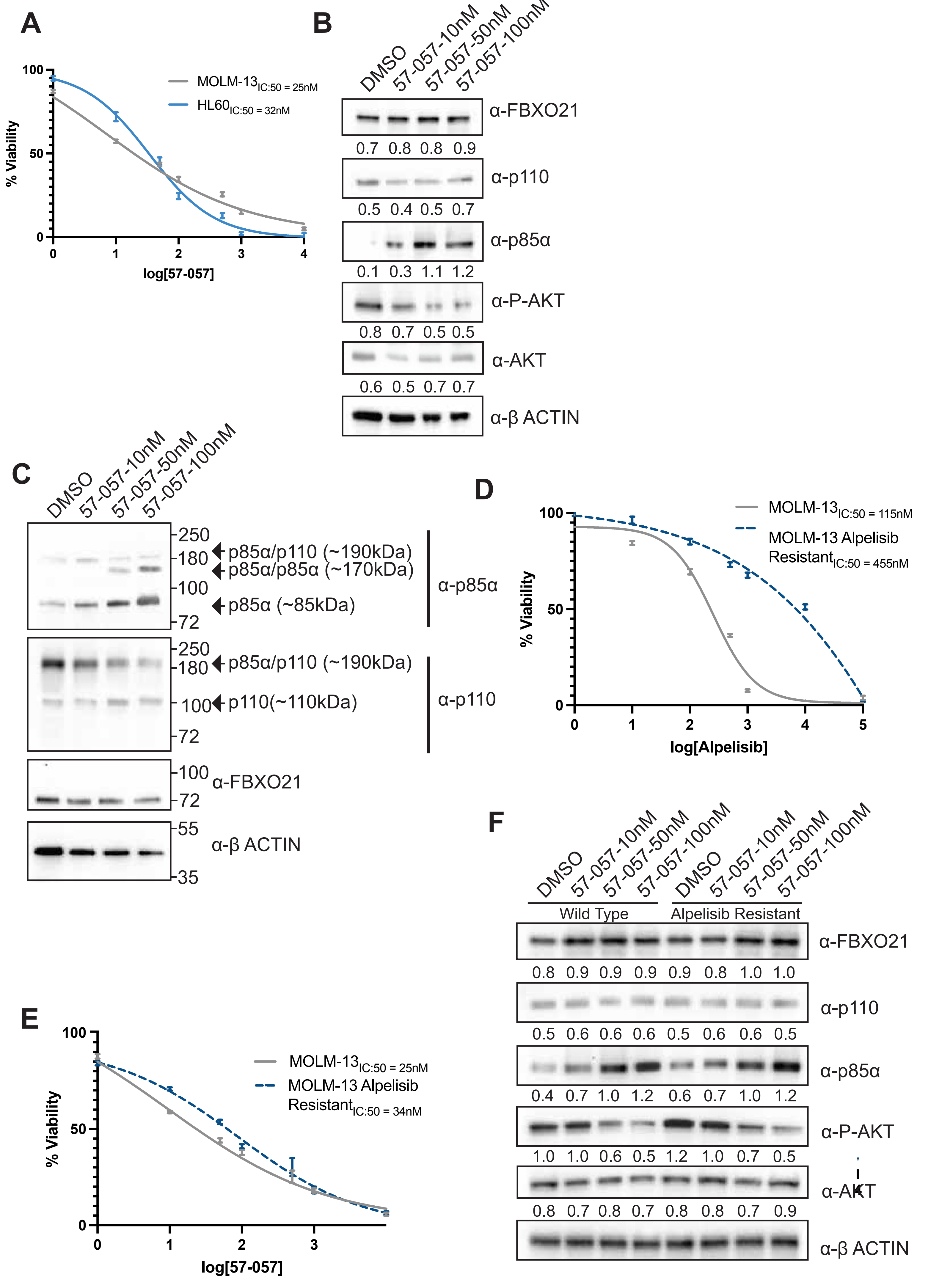

### Supplementary Figure 2

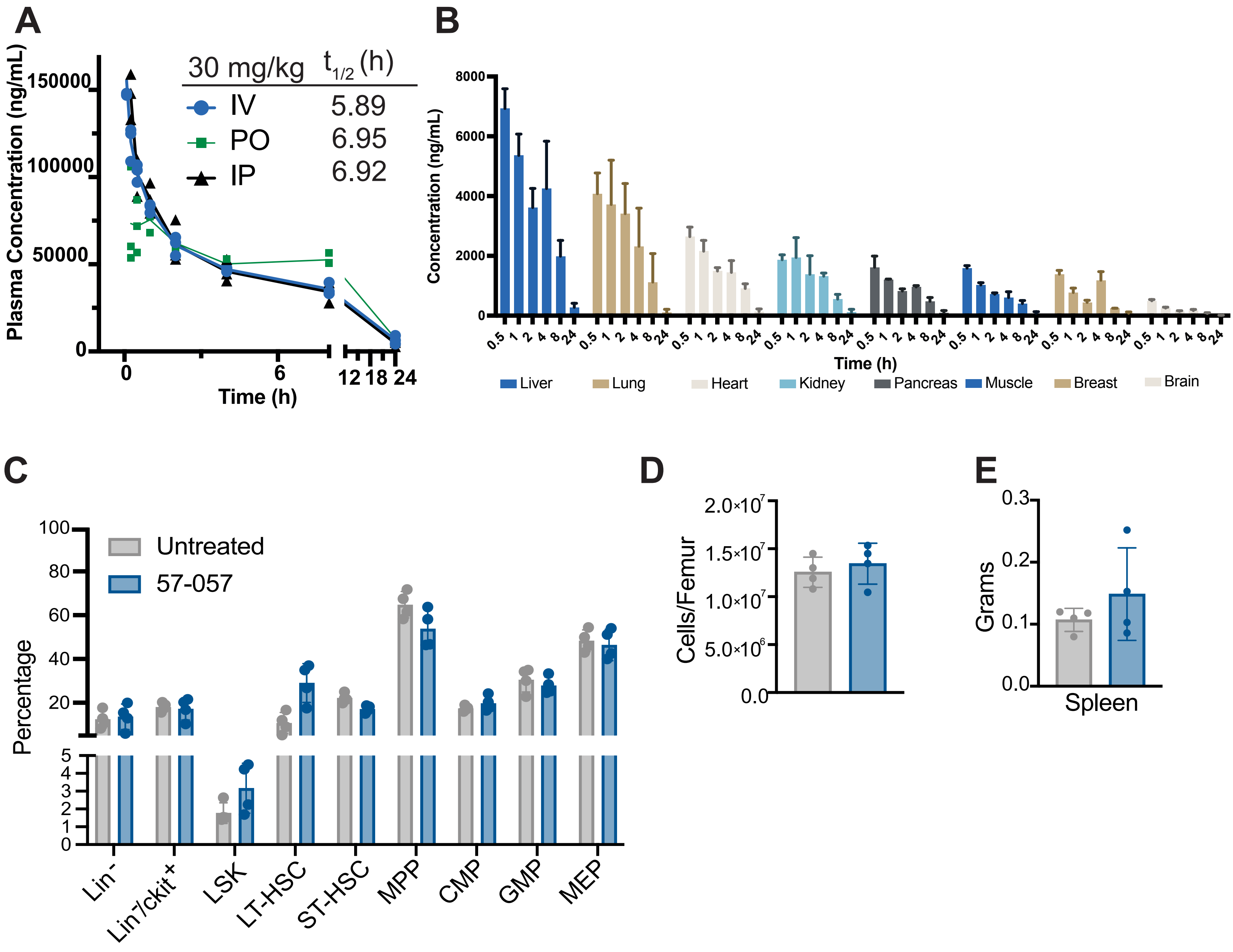

Supplemental Material

**Supplemental Figure 1: Sensitivity of AML cells to FBXO21 inhibitor 57-057 and its effect on PI3K signaling.** (n=3 for all experiments) **A** MTT assays for MOLM-13 and HL-60 cells treated with varying concentrations between 0.1 nM – 1 µM of 57-057 for 72 hours. **B** Western blot analysis of MOLM-13 cells treated with 57-057 at 10 nM, 50 nM, and 100 nM for 72 hours. The blot shows levels of FBXO21, p110, p85α, phosphorylated AKT (P-AKT), total AKT, and β-ACTIN as a loading control. **C** Native PAGE analysis of PI3K complex proteins in MOLM-13 cells treated with 57-057 at the indicated concentrations for 72 hours. The blot shows levels of p85α/p110 heterodimers, p85α/p85α homodimers, p85α monomers, p110, FBXO21, and β-ACTIN. **D** MTT assay for wild type MOLM-13 cells and MOLM-13 cells made resistant to Alpelisib (a PI3K inhibitor) treated with varying concentrations of Alpelisib between 1 nM – 10 µM for 72 hours. **E** MTT assay for MOLM-13 and Alpelisib-resistant MOLM-13 cells treated with 57-057 for 72 hours. **F** Western blot analysis of wild-type and Alpelisib-resistant MOLM-13 cells treated with 57-057 at 10 nM, 50 nM, and 100 nM for 72 hours. The blot shows levels of FBXO21, p110, p85α, phosphorylated AKT (P-AKT), total AKT, and β-ACTIN. Densitometry values are provided for each protein relative to β-ACTIN.

**Supplemental Figure 2: Pharmacokinetics and tissue distribution of 57-057 A** Plasma levels of 57-057 following IP, IV, and PO dosing at 30 mg/kg in mice. The half-lives ($t_1/_2$) for IV, PO, and IP routes are indicated. **B** Distribution of 57-057 in various tissues following a single IV administration (30 mg/kg). (n = 3). Dose escalation studies were conducted in mice treated with 57-057, starting at 2.5 mg/kg and increasing up to 90 mg/kg over a 14-week period. At sacrifice, one week following last dose, **C** bone marrow was analyzed for percentage of long-term hematopoietic stem cells (LT-HSCs), short-term hematopoietic stem cells (ST-HSCs), multipotent progenitors (MPPs), common myeloid progenitors (CMPs), granulocyte-monocyte progenitors (GMPs), and megakaryocyte-erythroid progenitors (MEPs). **D** Bone marrow cellularity and **E** spleen weight were also measured. (*p ≤ 0.05, **p ≤ 0.01, ***p ≤ 0.001)

**Table 1**

**Western Blot Antibodies:**

| Antibody | Company | Ref# | Dilution |
|---|---|---|---|
| AKT | Cell Signaling | 9272 | 1:1000 |
| Phospho-AKT | Cell Signaling | 4060 | 1:1000 |
| β-actin-HRP | Santa Cruz | sc-47778 | 1:10000 |
| FBXO21 | Abcam | [EPR13163] (ab179818) | 1:5000 |
| FLAG-HRP | Sigma | A8592 | 1:10000 |
| GFP | Cell Signaling | 2555 | 1:1000 |
| HA-HRP | Cell Signaling | 14031 | 1:10000 |
| mTOR | Cell Signaling | 2983 | 1:5000 |
| Phospho-mTOR (Ser2448) | Cell Signaling | 5536 | 1:5000 |
| c-Myc/n-Myc | Cell Signaling | 13987 | 1:1000 |
| p38 MAPK | Cell Signaling | 8690 | 1:5000 |
| Phospho-p38 MAPK (T180/Y182) | Cell Signaling | 4511 | 1:5000 |
| PI3 Kinase p85 | Cell Signaling | 4257 | 1:1000 |
| PI3 Kinase p85b | Abcam | [EPR18416] (ab180967) | 1:1000 |
| Phospho-PI3 Kinase p85 | Cell Signaling | 4228 | 1:1000 |
| p85α | Abcam | M253 | 1:1000 |
| PI3 Kinase p110α | Cell Signaling | 4255 | 1:1000 |
| PI3 Kinase p110b | Cell Signaling | C33D4 | 1:1000 |
| PTEN | Cell Signaling | 9552 | 1:1000 |
| Ubiquitin-HRP | Cell Signaling | 14049 | 1:10000 |
| K48 Ubiquitin | Cell Signaling | 8081 | 1:10000 |

**Table 2:** Addgene Plasmids

| Plasmid Name | Addgene ID |
|---|---|
| PI3KR1 human GFP tagged | 13515 |
| pRK5-HA-Ubiquitin-WT | 17608 |
| pRK5-HA-Ubiquitin-K6 | 22900 |
| pRK5-HA-Ubiquitin-K11 | 121152 |
| pRK5-HA-Ubiquitin-K27 | 17605 |
| pRK5-HA-Ubiquitin-K29 | 17606 |
| pRK5-HA-Ubiquitin-K48 | 17602 |
| pRK5-HA-Ubiquitin-K48R | 12201 |
| pRK5-HA-Ubiquitin-K33 | 17607 |

| | |
|---|---|
| pRK5-HA-Ubiquitin-K63 | 17604 |

**Table 3:** Cloning Primers

| Name | Sequence 5' to 3' |
|---|---|
| FBXO21_dYccV__F | CACTACATCCCAAATGCAG |
| FBXO21_dYccV__R | CAGACATCCCTGTGCTTC |
| p85_K692R_F | ACAGCTCTCTGCGCGAACTGGTGCTAC |
| p85_K692R_R | ACAAGTTATAGGGCTCGG |
| p85_K674R_F | TGTCATAAACAGAACAGCAACTG |
| p85_K674_R | CAATGCTTTACTTCGCCG |
| p85_K633R_F | CAACCGAAACAGAGCTGAAAACC |
| p85_K633R_R | CTGCTTCCAACATTCCATG |
| p85_K513,519R_F | CAATGAGAGAGAAATACAAAGGATTATGCATAATTATG |
| p85_K513,519R_R | CCTTCACGTCTAAACTTTTCTATGTATTCTTTGC |
| p85_K506R_F | GCGGTACAGCAGAGAATACATAG |
| p85_K506R_R | TCTTGGGTCTGGCACTGT |
| p85_K459R_F | GTTTCAAGAAAGAAGTCGAGAATATG |
| p85_K459R_R | TGAGTGTTATATTCATGTAATTTTTTC |
| p85_K448R_F | TGTAGGGAAAAGATTACATGAATATAAC |
| p85_K448R_R | GCTTCAATATTATCTTCTTTGAC |
| p85_K438R_F | TCAAGTTGTCAGAGAAGATAATATTG |
| p85_K438R_R | TCCTGTTGGTATTTGGATAC |
| p85_K419R_F | GTATAATCCCAGATTGGATGTGAAATTAC |
| p85_K419R_R | TGAGCTAGAGATTCATTCC |
| p85_K382R_F | CAAATTAATCAGAATATTTCATCGAGATG |
| p85_K382R_R | TTATTTCCCCCTTTCCTTAG |
| p85_dSH3_F | AAAAAAATCTCGCCTCCC |
| p85_dSH3_R | CATTCATCGCCTCTGCTG |
| p85_dSH3RhoGAP_F | CAGCCTGCACCAGCACTG |
| p85_dSH3RhoGAP_R | CATTCATCGCCTCTGCTGTG |
| p85_dSH2C_F | TATGCACAGCAGAGGCGA |
| p85_dSH2C_R | CATGTCTTCTCATCATGATGGG |
| p85_SH2C_F | AATGTTGGAAGCAGCAAC |
| p85_SH2C_R | CATTCATCGCCTCTGCTG |
| p85_SH2N_F | AAAAAAATCTCGCCTCCC |
| p85_SH2N_R | CATTCATCGCCTCTGCTG |
| p85_dSH2N_F | TCCAAATACCAACAGGATC |
| p85_dSH2N_R | CATTCAGCATCTTGTAAGG |
| p85_dSH2i_F | AATGTTGGAAGCAGCAAC |
| p85_dSH2i_R | ACTGGATAAAGTAATTTCACATC |
| p85_Y452A_F | ATTACATGAAGCTAACACTCAGTTTCAAGAAAAAAG |
| p85_Y452A_R | TTTTTCCCTACAGCTTCAATATTATC |
| p85_Y467A_F | TGATAGATTAGCTGAAGAATATACCCGC |
| p85_Y467A_R | TATTCTCGACTTTTTTCTTGAAAC |
| p85_Y580A_F | GAGAGACCAAGCTTTGATGTGGTTGAC |
| p85_Y580A_R | GTCTTTCTCAGCTGGATAAG |
| p85_Y607A_F | TGAAGACCAAGCTTCACTGGTGG |
| p85_Y607A_R | GTGTTTTCATTGCCCAAC |

**Table 4:** Flow Cytometry Antibodies

| Antibody | Fluorochrome | Clone | Company |
|---|---|---|---|
| Annexin V | APC | n/a | BioLegend |
| Propodium Iodide (PI) | | n/a | BioLegend |

**Viral transduction and additional cell culture**

Lentivirus was made through Lipofectamine 2000 transfection of HEK293T cells, with shRNA plasmid, Pax2, and pMD2.g plasmids, and retrovirus was made through Lipofectamine 2000 transfection of Phoenix-AMPHO cells with pMIG plasmid. Transfection media was changed after 24 hours, and viral media was collected and concentrated after 48 hours. AML cell lines and patient samples were transduced with virus for 1 hour at 1000xg and 37°C. 1-4 million cells were resuspended in 500µL media in a 15mL conical tube with 100µL concentrated virus and 1µL polybrene (10mg/mL). Cells were then resuspended in viral media, plated in 24 well plate, and left to rest for 3 hours at 37°C. After 3 hour rest, cells were resuspended in media and treated with 1µg/ml of puromycin 48 hours post infection or sorted using fluorescent selection marker. AML primary cells were thawed and cultured overnight prior to viral transduction.

**Western blot analysis and immunoprecipitation**

For western blot analysis, samples were lysed in lysis buffer (20mM Tris pH7.5, 150mM NaCl, 1mM EDTA) containing 1X Halt Protease and Phosphatase Inhibitor Cocktail (ThermoFisher, Waltham, MA, USA) and 10 mM N-ethylmaleimide. For native gel western blot analysis, cells were lysed as described above and loaded with 2x sample buffer (62.5mM Tris-HCl, pH 6.8, 40% glycerol, 1% bromophenol blue) and Tris/Glycine running buffer (25mM Tris, 192mM glycine, pH 8.3). Antibodies are listed in supplemental materials and methods.

Quantification is reflective of relative fold change as a ratio of each protein band relative to our loading control, β-actin utilizing relative densitometry calculated with ImageJ Software.

**Flow Cytometry Analysis**

For apoptosis, staining was performed following BioLegend apoptosis staining protocol. Antibodies listed in supplemental materials and methods.

**General Chemistry Experimental Methods**

Unless otherwise stated, all the reactions were performed in oven- or flame-dried glassware under an argon atmosphere. Air and/or moisture-sensitive liquids were transferred via a syringe or stainless-steel cannula. Reactions were examined by thin-layer chromatography (TLC) on glass plates pre-coated with silica gel (0.25 mm, 60 Å pore size, 230–400 mesh, Merck KGA) impregnated with a fluorescent indicator (254 nm), visualized under UV light (254 nm). Organic solutions were concentrated under reduced pressure on a Buchi rotary evaporator. Flash chromatography was performed using Buchi Pure C-850 (230-400 mesh, 40-63 μm prepacked SiliCycle silica cartridges).

**Chemistry Materials and Reagents**

All commercially available reagents were purchased from Sigma-Aldrich, Merck or Ambeed and used without purification unless otherwise indicated. All organic solvents used in the experiment were anhydrous. Extraction and chromatography solvents were reagent grade and used without purification.

**Instrumentation**

Proton nuclear magnetic resonance ($^{1}$HNMR) spectra, proton-decoupled carbon nuclear magnetic resonance ($^{13}$CNMR) spectra, and proton-decoupled fluorine nuclear magnetic resonance ($^{19}$FNMR) spectra were recorded on Bruker Advance 400 (400 MHz) and 600 (600 MHz) spectrometers using $CDCl_3$ and DMSO-$d_6$ as the solvents and tetramethylsilane (TMS) as internal standard at room temperature. All chemical shifts (δ) are reported in parts per million (ppm) downfield from tetramethylsilane and are referenced to the residual solvent signal of the

NMR solvent. Data are represented as follows: chemical shift, multiplicity (bs = broad singlet, s = singlet, d = doublet, t = triplet, q = quartet, m = multiplet), coupling constants (*J*) in Hertz (Hz), integration. High-Resolution Mass Spectrometry (HRMS) was performed on an Agilent 6230 LC/TOF mass spectrometer with an ESI source running on a Poroshell 120, EC-C18 column (4.6 × 50 mm, 2.7 µm) using water with 0.1% formic acid as solvent A and acetonitrile with 0.1% formic acid as solvent B. The gradient for elution varied from 30% B to 95% B over 7 minutes at a flow rate of 0.5 mL/min. High-Resolution Gas Chromatography-Mass Spectrometry (HR-GCMS) analysis was performed on an Agilent 7250 GC/Q-TOF mass spectrometer coupled with an Agilent 8890 GC. Separation was achieved on an HP-5ms Ultra Inert column (30 m × 250 µm × 0.25 µm). The oven temperature was programmed to increase at a rate of 10°C/minute, reaching a final temperature of 300°C. Helium was used as the carrier gas, with a flow rate of 1 mL/minute. . The purity of all the compounds was confirmed to be >95% by High Performance Liquid Chromatography (HPLC) on an Agilent 1260 Infinity instrument running on an Eclipse plus C8 column (4.6 x 150 mm, 3.5 µm) with an attached DAD detector. For analytical HPLC, the gradient for elution varied from 30% B to 95% B over 8 minutes, followed by an isocratic elution for 4 minutes at a flow rate of 0.6 mL/min.

**Synthesis**

**General scheme for the synthesis of terphenyl derivatives**

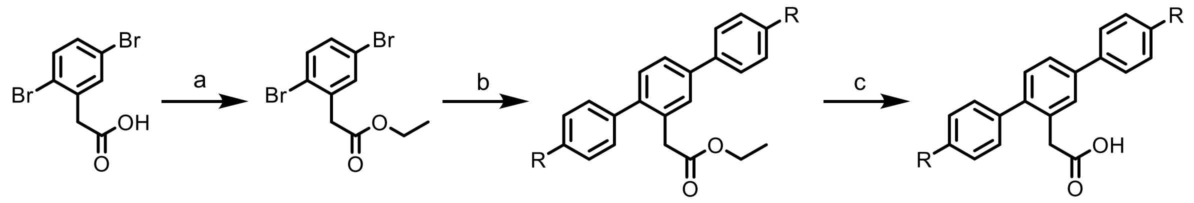


**1** **2** **3** **4**

**R = F; 57-055**
**R = OH; 57-048**
**R = CN; 57-119**

**R = F; 57-057**
**R = OH; 57-112**
**R = CN; 57-125**

Reaction conditions: (a) Anhydrous. EtOH, cat. conc. $H_2SO_4$ (0.2 eq.), reflux, 5h; (b) R-PhB(OH)$_2$ (3 eq.), $Na_2CO_3$ (3 eq.), Pd(OAc)$_2$ (0.1 eq.), $H_2O$: Dioxane (1:2), 80 °C, 12h; (c) MeOH, cat. 1N NaOH, 70 °C, 2h.

**Synthesis of Ethyl 2-(2,5-dibromophenyl)acetate (2)[1]**

To a stirred solution of **1** (1.0 equiv) in anhydrous ethanol (10 mL) in an oven-dried round bottom flask was added a catalytic amount of concentrated $H_2SO_4$ (0.2 eq.). The round bottom flask was placed in a preheated oil bath and refluxed at 80 °C for 5h. Upon completion (monitored by TLC), the reaction mixture was cooled to room temperature and concentrated under reduced pressure. The residue was neutralized with an aqueous $NaHCO_3$ solution. The precipitate was filtered, washed with water (3 × 5 mL), and dried under a high vacuum to give **2** as a buff-colored solid. The crude was characterized and used for **step b** without further purification.

**General Method for the Synthesis of 3 using Suzuki-Miyaura Cross-Coupling Reaction**

An oven-dried sealed tube was charged with **2** (1.0 equiv), substituted phenylboronic acid (3.0 equiv), and $Na_2CO_3$ (3.0 equiv). The reaction tube was evacuated and backfilled with argon (this process was repeated a total of three times). Anhydrous dioxane (4 mL) and $H_2O$ (2 mL) were added through a syringe. The resulting reaction mixture was purged with argon for 10 minutes. $Pd(OAc)_2$ (0.1 equiv) was then added, and the sealed tube was capped and placed in a preheated oil bath and stirred at 80 °C for 12h. Upon completion (monitored by TLC), the reaction mixture was cooled to room temperature, poured into cold water, and extracted with EtOAc (3 × 10 mL). The combined organic layer was washed with brine, dried over a $Na_2SO_4$ bed, and concentrated under reduced pressure. The crude residue was purified by flash chromatography (2-10% EtOAc in Hexane) to yield **3**.

**General Method for the Synthesis of 4 using Base-catalyzed Hydrolysis of Esters**

To a stirred solution of **3** (1.0 equiv) in methanol (5 mL) in an oven-dried round bottom flask, a catalytic amount of 1N NaOH (0.1 mL) was added. The resulting reaction mixture was placed in a preheated oil bath and refluxed at 70 °C for 2h. Upon completion (monitored by TLC), the reaction mixture was cooled to room temperature and concentrated under reduced pressure. The residue was acidified with 1N HCl solution. The precipitate was filtered, washed with water (3 × 5 mL), and dried under a high vacuum**.** The crude residue was purified by flash chromatography

(50-70% EtOAc in Hexane) to yield **4**.

**Synthesis of 2'-ethyl-4,4''-difluoro-1,1':4',1''-terphenyl (57-139)**

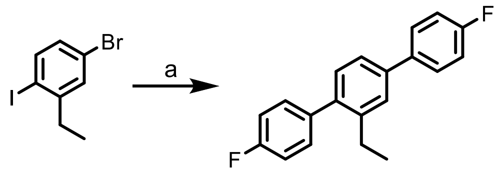


**2** **57-139**

Reaction conditions: (a) 4F-PhB$(OH)_2$ (2.3 eq.), $Na_2CO_3$ (3 eq.), $Pd(PPh_3)_4$ (0.05 eq.), $H_2O$: Dioxane (1:2), 80 °C, 12h (54%).

An oven-dried sealed tube was charged with the **2** (1.0 equiv), 4-fluoro phenylboronic acid (2.3 equiv), and $Na_2CO_3$ (3.0 equiv). The reaction tube was evacuated and backfilled with argon (this process was repeated a total of three times). Anhydrous dioxane (4 mL) and $H_2O$ (2 mL) were then added through a syringe. The resulting reaction mixture was purged with argon for 10 minutes. $Pd(PPh_3)_4$ (0.05 equiv) was then added, and the sealed tube was capped and placed in a preheated oil bath and stirred at 80 °C for 12h. Upon completion (monitored by TLC), the reaction mixture was cooled to room temperature, poured into cold water, and extracted with EtOAc (3 × 10 mL). The combined organic layer was washed with brine, dried over a $Na_2SO_4$ bed, and concentrated under reduced pressure. The crude residue was purified by flash chromatography (1% EtOAc in Hexane) to yield **57-139** as a white solid.

**Characterization of compounds**

**Ethyl 2-(2,5-dibromophenyl)acetate (2):**

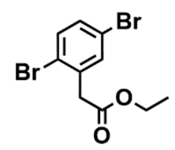


Buff solid, 520.0 mg, yield 95%. $^{1}$HNMR (600 MHz, $CDCl_3$) δ 7.43 – 7.41 (m, 2H), 7.26 (dd, $J$ = 8.6, 2.5 Hz, 1H), 4.19 (q, 2H), 3.73 (s, 2H), 1.27 (t, 3H). $^{13}$CNMR (151 MHz, $CDCl_3$) δ 169.83, 136.35, 134.28, 134.09, 131.87, 123.73, 121.22, 61.27, 41.47, 14.17. HRMS (ESI-TOF) m/z: $[M+H]^+$ Calcd for $C_{10}H_{10}Br_2O_2$ 320.9126; found 320.9128.

**Ethyl 2-(4,4''-difluoro-[1,1':4',1''-terphenyl]-2'-yl)acetate (57-055):**

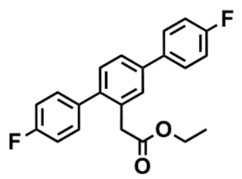

White solid, 108.7 mg, yield 66%. [1]HNMR (600 MHz, $CDCl_3$) δ 7.60 – 7.57 (m, 2H), 7.53 (d, *J* = 1.8 Hz, 1H), 7.50 (dd, *J* = 7.9, 1.9 Hz, 1H), 7.33 – 7.31 (m, 3H), 7.15 – 7.10 (m, 4H), 4.10 (q, 2H), 3.62 (s, 2H), 1.21 (t, 3H). [13]CNMR (151 MHz, CDCl3) δ 171.70, 162.57 (d, *J* = 247.8 Hz), 162.23 (d, *J* = 245.1 Hz), 140.45, 139.66, 136.62 (d, *J* = 3.3 Hz), 136.60 (d, *J* = 2.8 Hz), 132.50, 130.89 (d, *J* = 8.3 Hz), 130.78, 129.08, 128.70 (d, *J* = 8.4 Hz), 125.80, 115.68 (d, *J* = 21.6 Hz), 115.18 (d, *J* = 21.9 Hz), 60.90, 39.06, 14.17. [19]FNMR (564 MHz, CDCl3) δ -115.32 (s), -115.46 (s). HRMS (ESI-TOF) m/z: $[M+H]^+$ Calcd for $C_{22}H_{18}F_2O_2$ 353.1353; found 353.1357.

**Ethyl 2-(4,4''-dihydroxy-[1,1':4',1''-terphenyl]-2'-yl)acetate (57-048):**

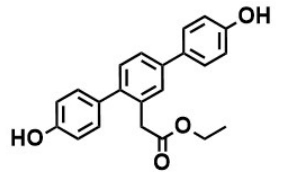

57-048 was synthesized using the procedure for **57-055**. Violet solid, 55.7 mg, yield 30%. [1]HNMR (400 MHz, DMSO-$d_6$) δ 9.60 (s, 1H), 9.55 (s, 1H), 7.52 – 7.49 (m, 4H), 7.22 (d, *J* = 7.9 Hz, 1H), 7.11 – 7.09 (m, 2H), 6.87 (d, *J* = 8.4 Hz, 2H), 6.81 (d, *J* = 9.1 Hz, 2H), 3.99 (q, 2H), 3.65 (s, 2H), 1.10 (t, 3H). [13]CNMR (151 MHz, DMSO-$d_6$) δ 170.10, 137.48, 136.36, 135.11, 134.64, 132.37, 127.46, 124.13, 120.98, 116.13, 116.05, 114.86, 113.69, 61.14, 40.99, 14.49. HRMS (ESI-TOF) m/z: $[M+H]^+$ Calcd for $C_{22}H_{20}O_4$ 349.1440; found 349.1435.

**Ethyl 2-(4,4''-dicyano-[1,1':4',1''-terphenyl]-2'-yl)acetate (57-119):**

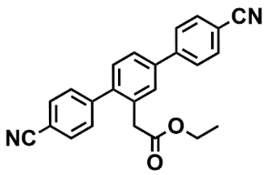

57-119 was synthesized using the procedure for **57-055**. White solid, 214.6 mg, yield 38%. [1]HNMR (400 MHz, CDCl3) δ 7.77 – 7.71 (m, 6H), 7.62 (d, *J* = 1.8 Hz, 1H), 7.59 (dd, *J* = 7.9, 1.9 Hz, 1H), 7.50 – 7.48 (m, 2H), 7.36 (d, *J* = 7.9 Hz, 1H), 4.11 (q, 2H), 3.62 (s, 2H), 1.21 (t, 3H). [13]CNMR (101 MHz, CDCl3) δ 171.19, 145.16, 144.60, 140.92, 139.42, 132.69, 132.60, 132.17, 130.66, 130.04, 129.66, 127.77, 126.27, 118.77, 118.65, 111.56, 111.41, 61.17, 38.88, 14.16. HRMS (ESI-TOF) m/z: $[M+H]^+$ Calcd for $C_{24}H_{18}N_2O_2$ 367.1447; found 367.1445.

**2-(4,4''-difluoro-[1,1':4',1''-terphenyl]-2'-yl)acetic acid (57-057):**

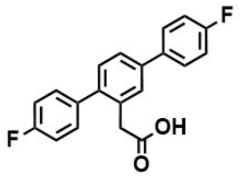

White solid, 44.0 mg, yield 95%. $^{1}$HNMR (400 MHz, DMSO-$d_6$) δ 12.30 (s, 1H), 7.75 (dd, $J$ = 8.7, 5.5 Hz, 2H), 7.67 – 7.59 (m, 2H), 7.40 – 7.26 (m, 7H), 3.60 (s, 2H). $^{13}$CNMR (101 MHz, DMSO-$d_6$) δ 173.17, 162.42 (d, $J$ = 244.4 Hz), 162.0 (d, $J$ = 244.1 Hz), 140.40, 138.68, 137.16 (d, $J$ = 3.4 Hz), 136.53 (d, $J$ = 2.8 Hz), 133.76, 131.34 (d, $J$ = 8.0 Hz), 130.94, 129.70, 129.07 (d, $J$ = 8.0 Hz), 125.62, 116.27 (d, $J$ = 21.6 Hz), 115.66 (d, $J$ = 21.4 Hz), 39.11. $^{19}$FNMR (564 MHz, DMSO-$d_6$) δ -115.31 (s), -115.38 (s). HRMS (ESI-TOF) m/z: $[M+H]^+$ Calcd for $C_{20}H_{14}F_2O_2$ 325.1040; found 325.1044.

**2-(4,4''-dihydroxy-[1,1':4',1''-terphenyl]-2'-yl)acetic acid (57-112):**

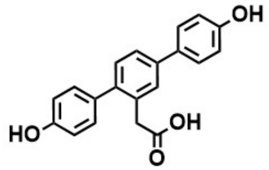

57-112 was synthesized using the procedure for **57-057**. White solid, 24.5 mg, yield 88%. $^{1}$HNMR (400 MHz, DMSO-$d_6$) δ 12.24 (bs, 1H), 9.54 (s, 1H), 9.50 (s, 1H), 7.52 – 7.47 (m, 4H), 7.21 (d, $J$ = 7.9 Hz, 1H), 7.12 (d, $J$ = 8.2 Hz, 2H), 6.86 (d, $J$ = 8.6 Hz, 2H), 6.82 (d, $J$ = 8.4 Hz, 2H), 3.56 (s, 2H). $^{13}$CNMR (101 MHz, DMSO-$d_6$) δ 173.45, 157.56, 156.99, 140.44, 139.03, 133.28, 131.57, 130.95, 130.75, 130.42, 128.82, 128.05, 124.81, 116.21, 115.53, 39.12. HRMS (ESI-TOF) m/z: $[M-H]^+$ Calcd for $C_{20}H_{16}O_4$ 319.0971; found 319.0975.

**2-(4,4''-dicyano-[1,1':4',1''-terphenyl]-2'-yl)acetic acid (57-125):**

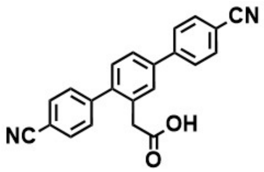

57-125 was synthesized using the procedure for **57-057**. White solid, 46.7 mg, yield 92%. $^{1}$HNMR (600 MHz, DMSO-$d_6$) δ 12.38 (s, 1H), 7.98 - 7.94 (m, 6H), 7.82 (d, $J$ = 1.9 Hz, 1H), 7.78 (dd, $J$ = 8.0, 2.0 Hz, 1H), 7.57 (d, $J$ = 8.4 Hz, 2H), 7.41 (d, $J$ = 8.0 Hz, 1H), 3.65 (s, 2H). $^{13}$CNMR (101 MHz, DMSO-$d_6$) δ 172.87, 145.56, 144.30, 141.09, 138.45, 133.95, 133.43, 132.80, 130.94, 130.41, 128.03, 126.25, 119.30, 119.24, 110.76, 39.02. HRMS (ESI-TOF) m/z: $[M+H]^+$ Calcd for $C_{22}H_{14}N_2O_2$ 339.1133; found 339.1134.

**2'-ethyl-4,4''-difluoro-1,1':4',1''-terphenyl (57-139):**

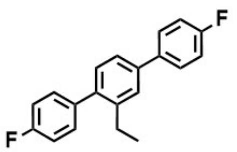

White solid, 206.1 mg, yield 54%. $^{1}$HNMR (400 MHz, $CDCl_3$) δ 7.64 – 7.59 (m, 2H), 7.50 (d, *J* = 1.9 Hz, 1H), 7.43 (dd, *J* = 7.9, 2.0 Hz, 1H), 7.35 – 7.31 (m, 2H), 7.28 (s, 1H), 7.19 – 7.12 (m, 4H), 2.67 (q, 2H), 1.17 (t, 3H). $^{13}$CNMR (151 MHz, CDCl3) δ 162.49 (d, *J* = 246.5 Hz), 162.01(d, *J* = 244.6 Hz), 142.22, 139.65, 139.60, 137.39 (d, *J* = 3.4 Hz), 137.10 (d, *J* = 3.3 Hz), 130.73 (d, *J* = 7.9 Hz), 130.58, 128.65 (d, *J* = 8.6 Hz), 127.35, 124.28, 115.63 (d, *J* = 21.1 Hz), 115.02 (d, *J* = 21.6 Hz), 26.29, 15.66. $^{19}$FNMR (564 MHz, $CDCl_3$) δ -115.71 (s), -115.89 (s). HR-GCMS m/z: $[M]^+$ for $C_{20}H_{16}F_2$: 294.1220; found 294.1207.

$^{1}$HNMR of **2** in $CDCl_3$

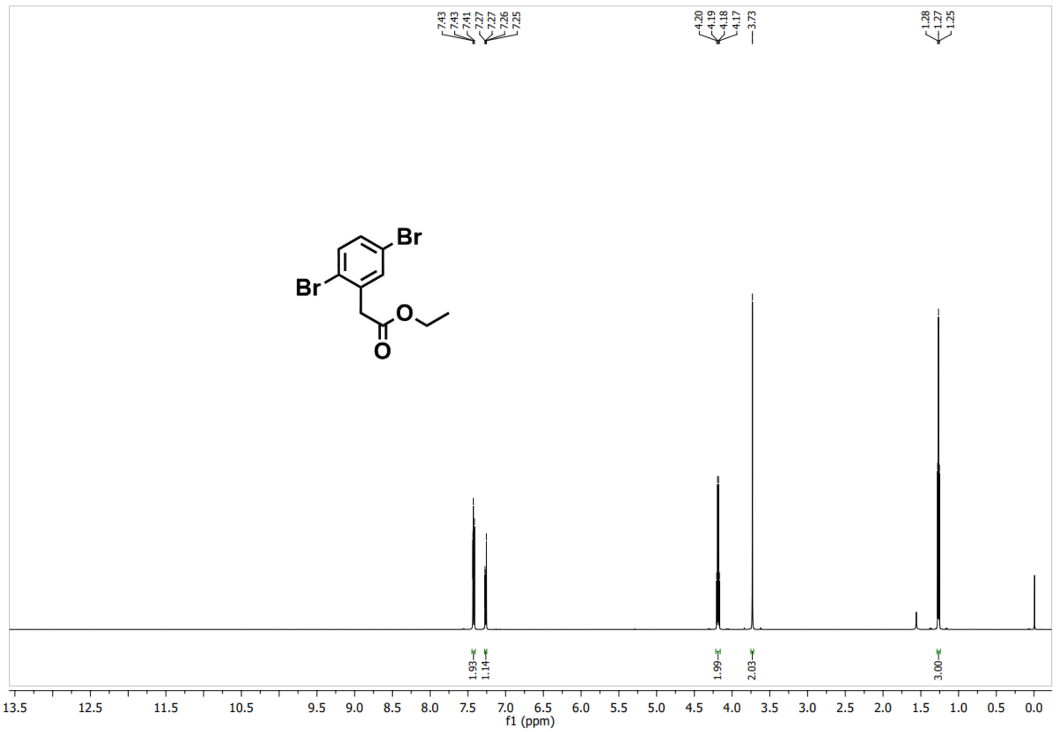


$^{13}$CNMR of **2** in $CDCl_3$

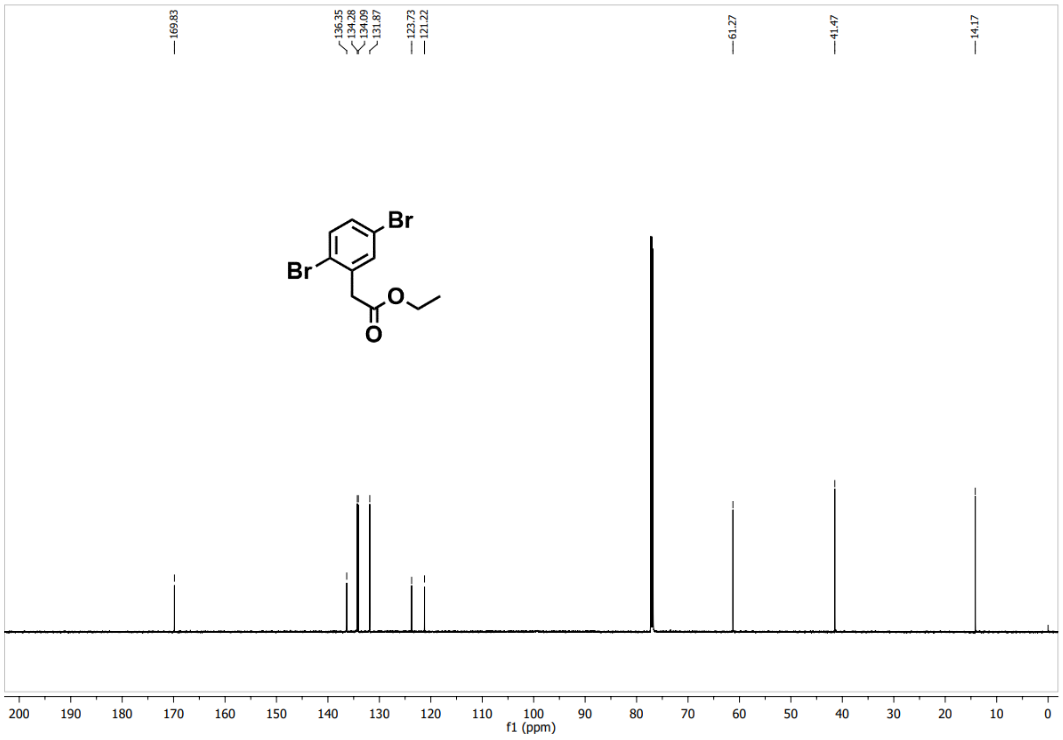


$^{1}$HNMR of **57-055** in $CDCl_3$

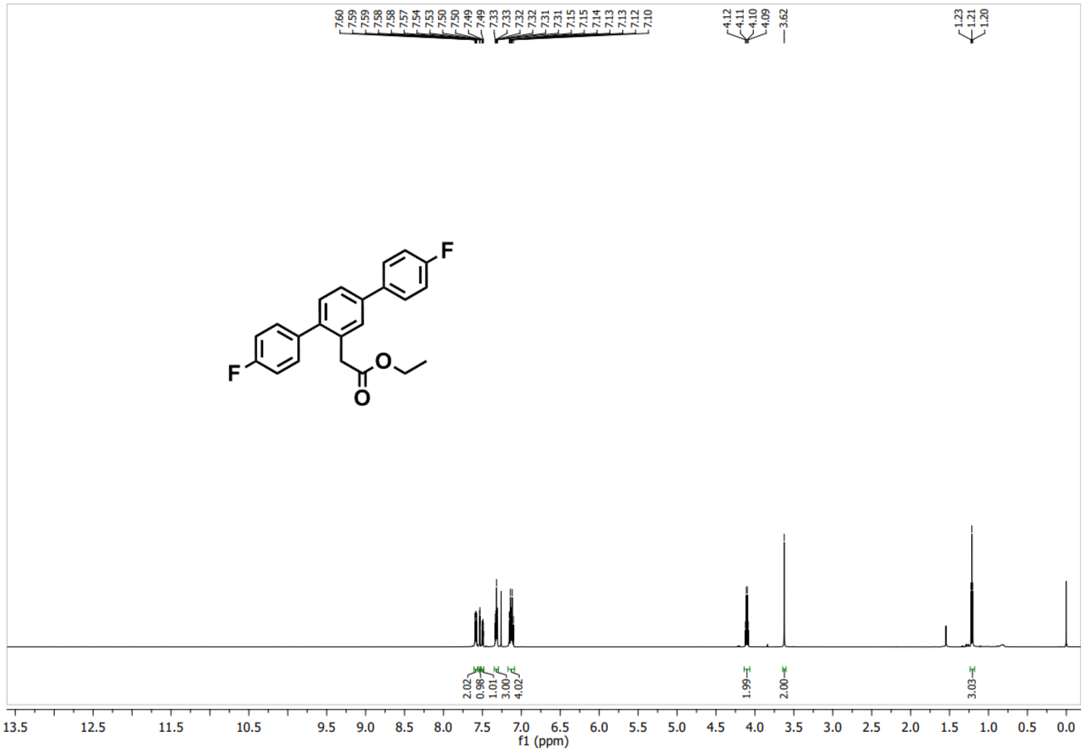


$^{13}$CNMR of **57-055** in $CDCl_3$

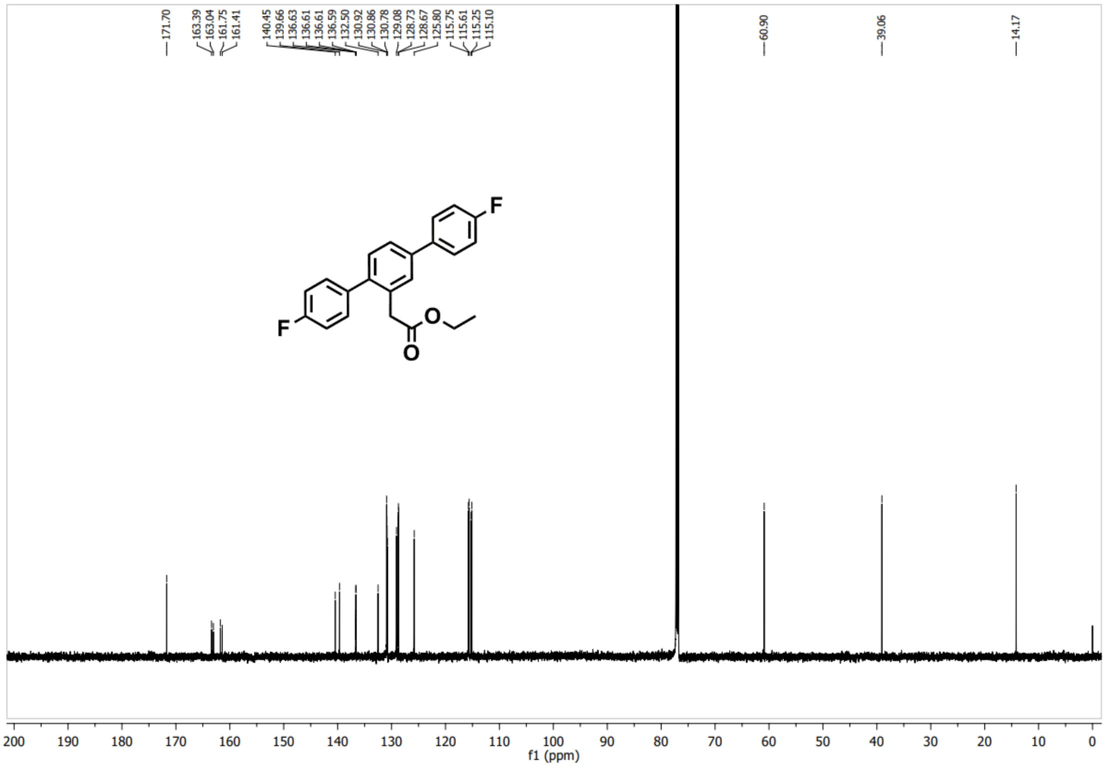


$^{19}$FNMR of **57-055** in $CDCl_3$

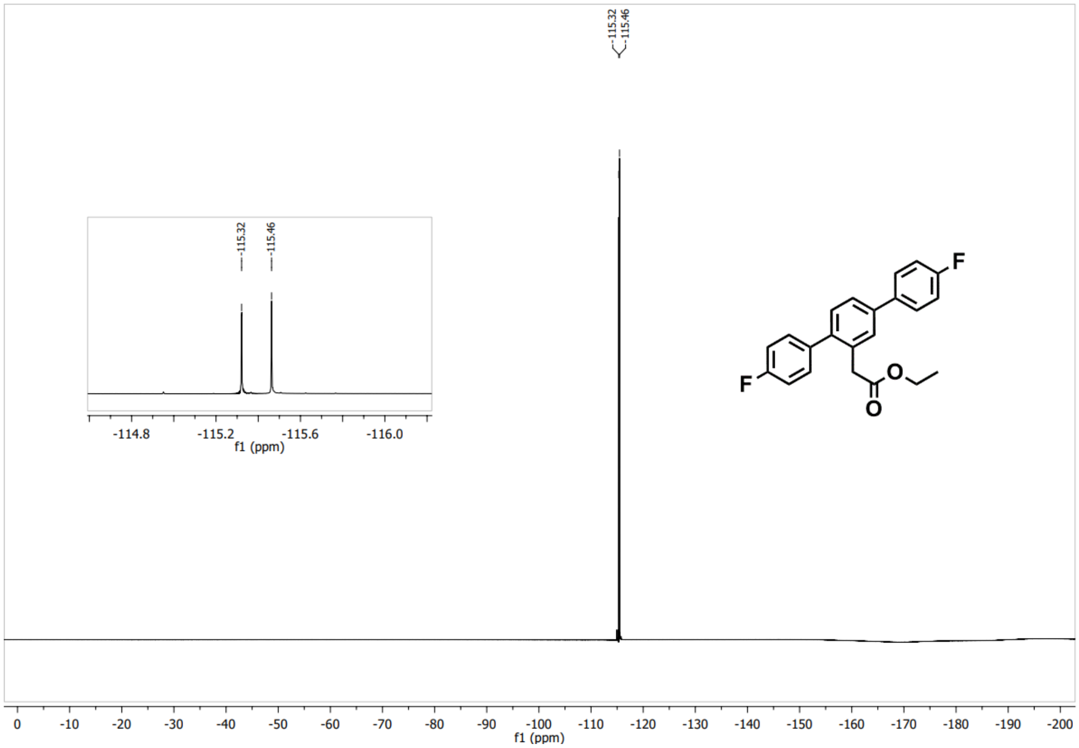

$^{1}$HNMR of **57-048** in DMSO-$d_6$

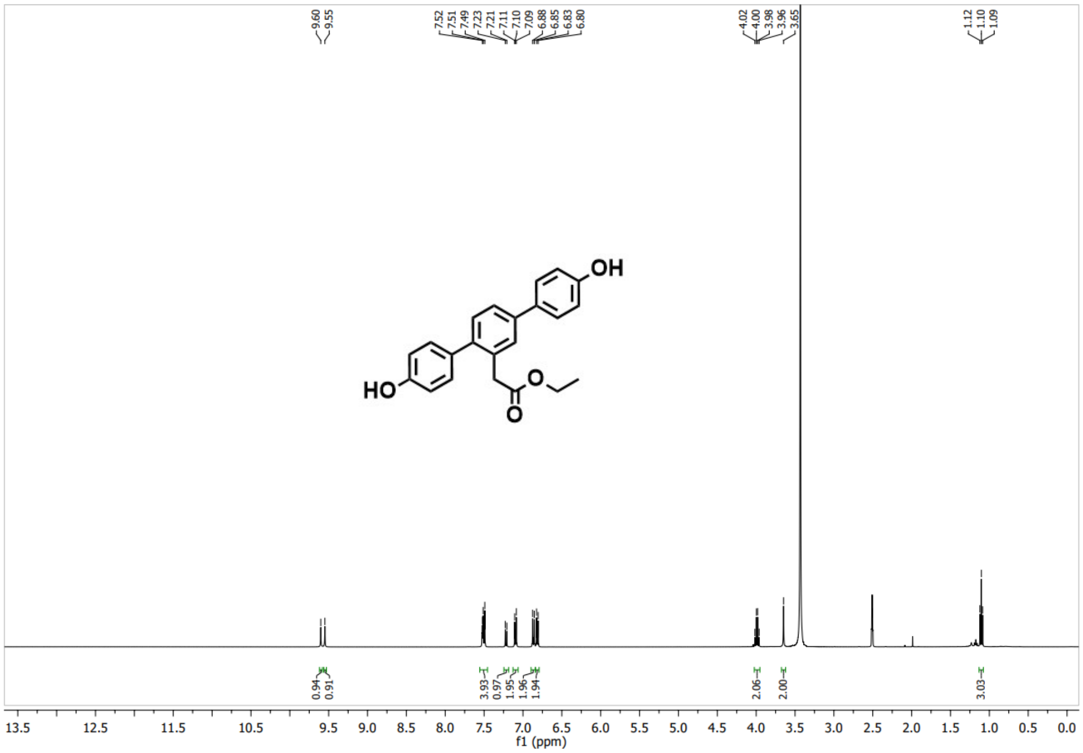

$^{13}$CNMR of **57-048** in DMSO-$d_6$

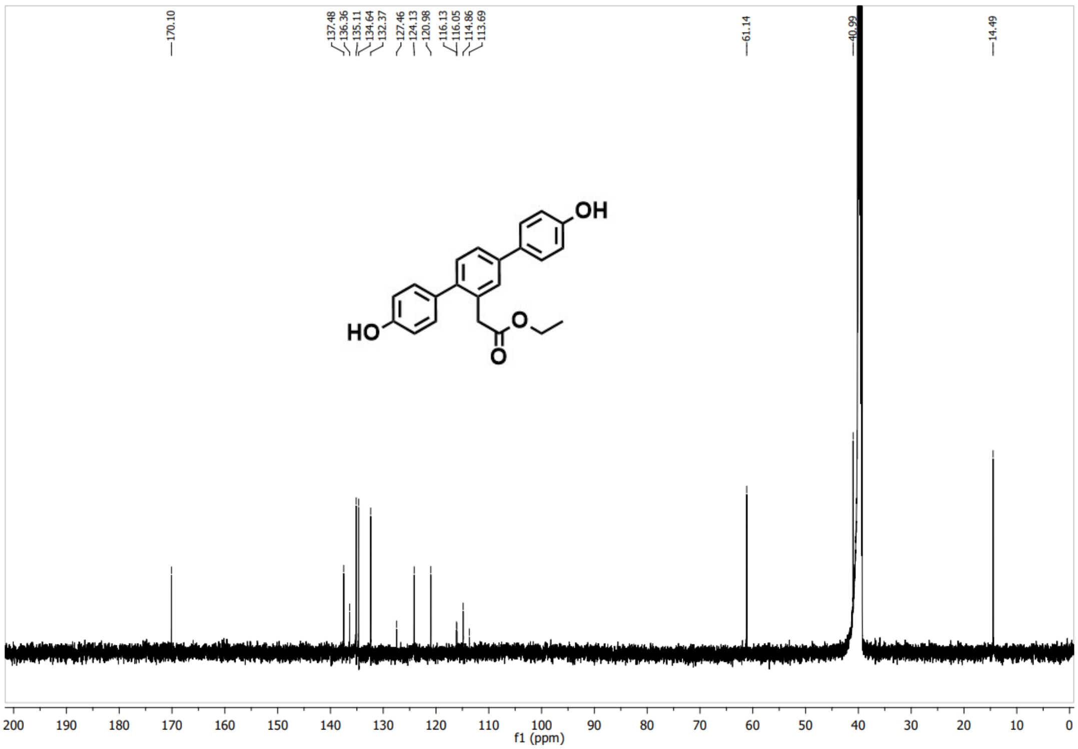


HPLC analysis of **57-048**

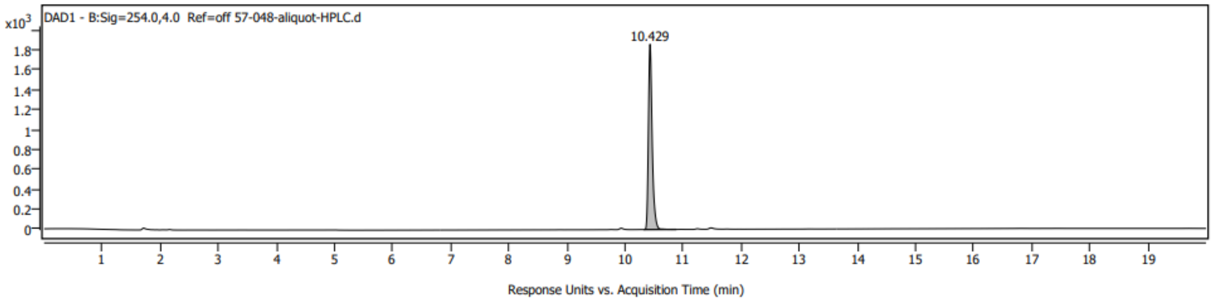


$^1$HNMR of **57-119** in $CDCl_3$

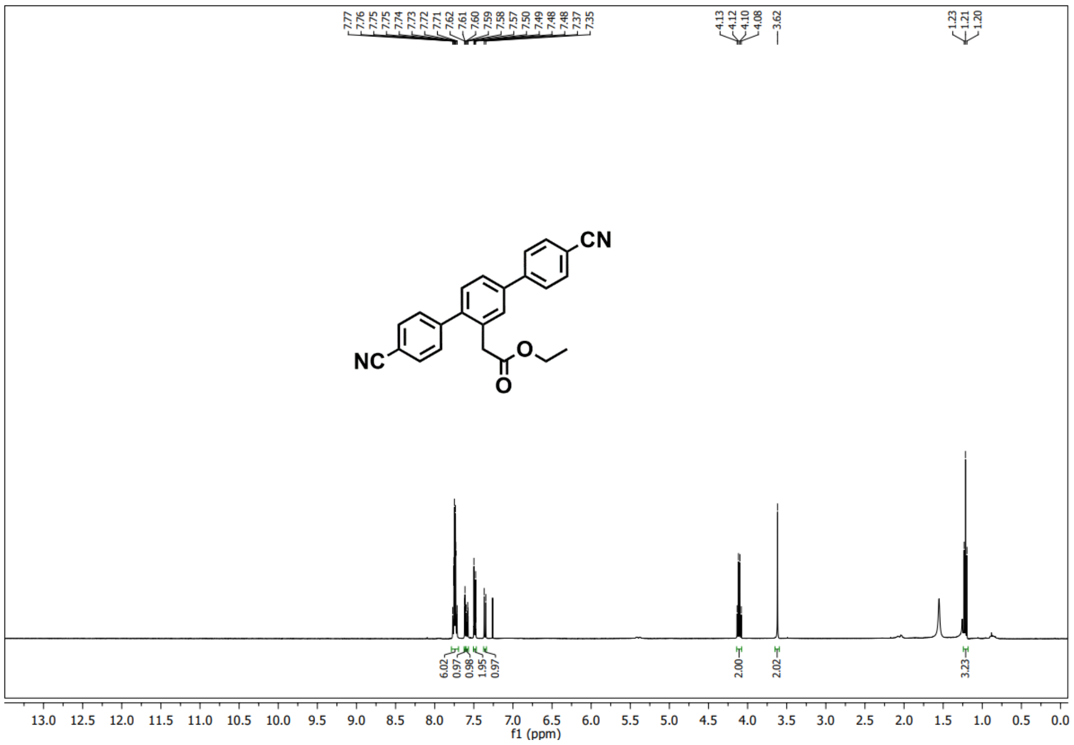


$^{13}$CNMR of **57-119** in $CDCl_3$

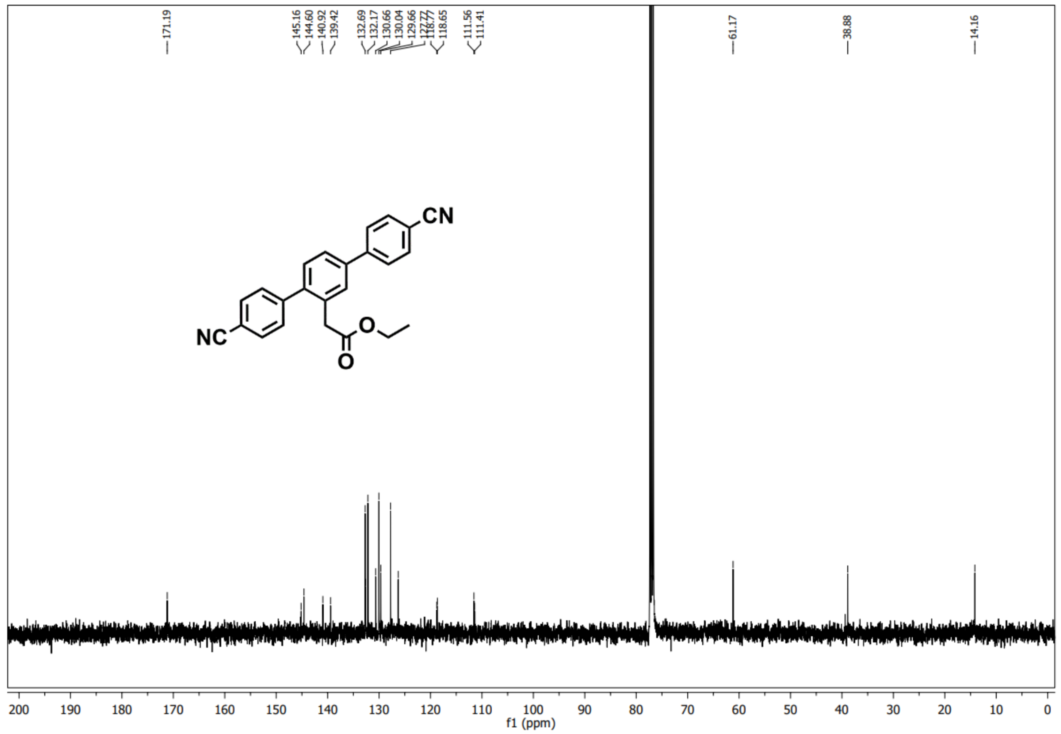


$^{1}$HNMR of **57-057** in DMSO-$d_6$

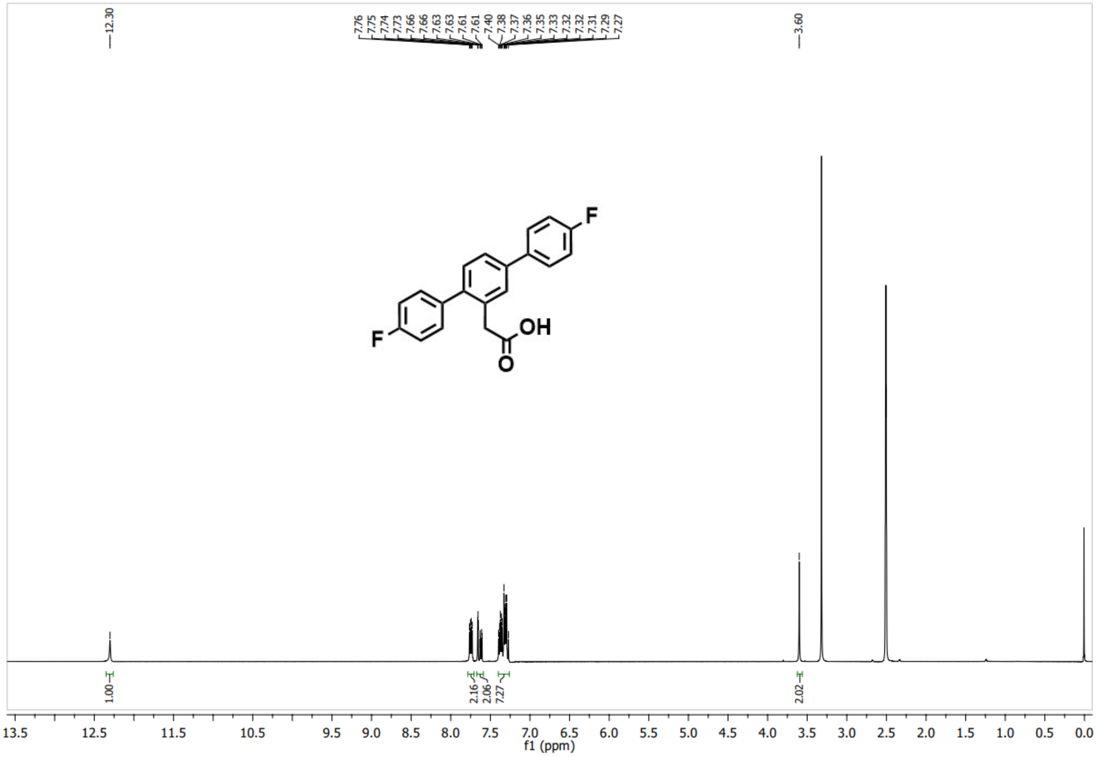

$^{13}$CNMR of **57-057** in DMSO-d$_6$

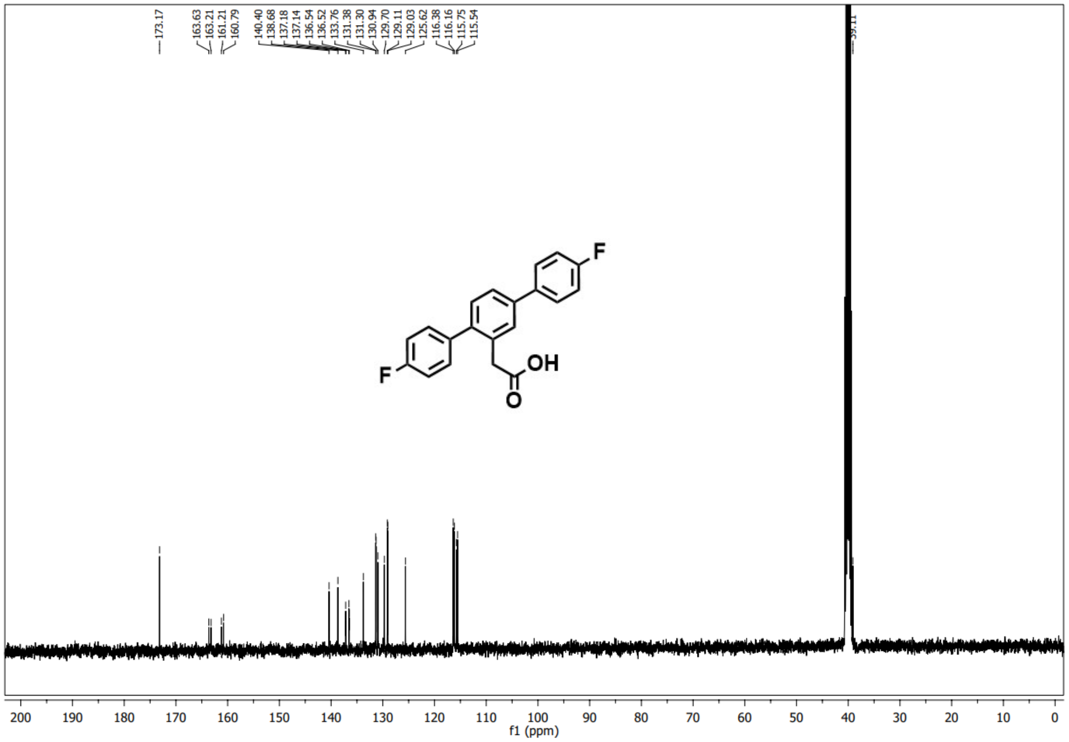

$^{19}$FNMR of **57-057** in DMSO-d$_6$

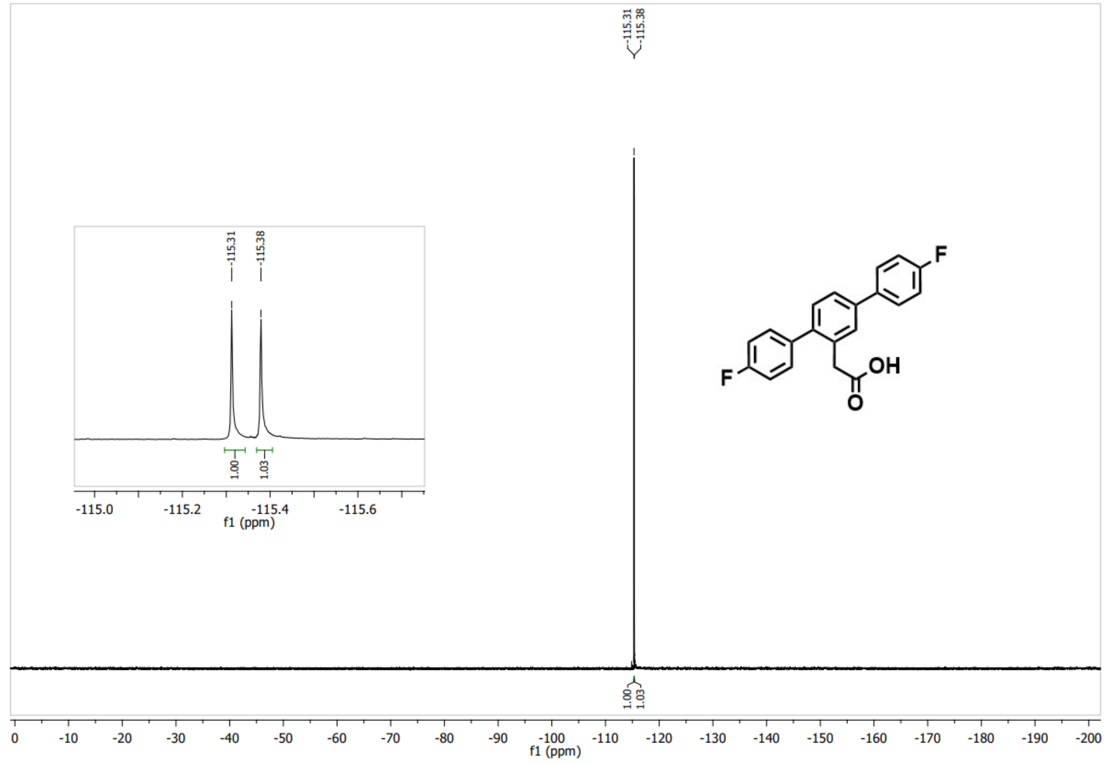


HPLC analysis of **57-057**

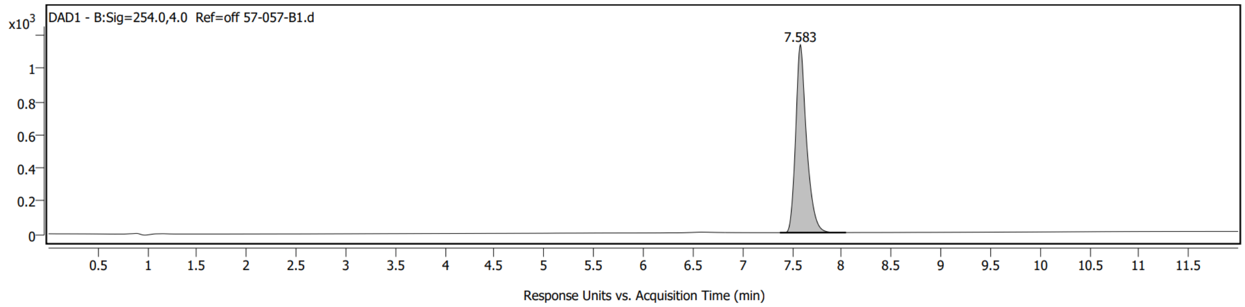


$^1$HNMR of **57-112** in DMSO-$d_6$

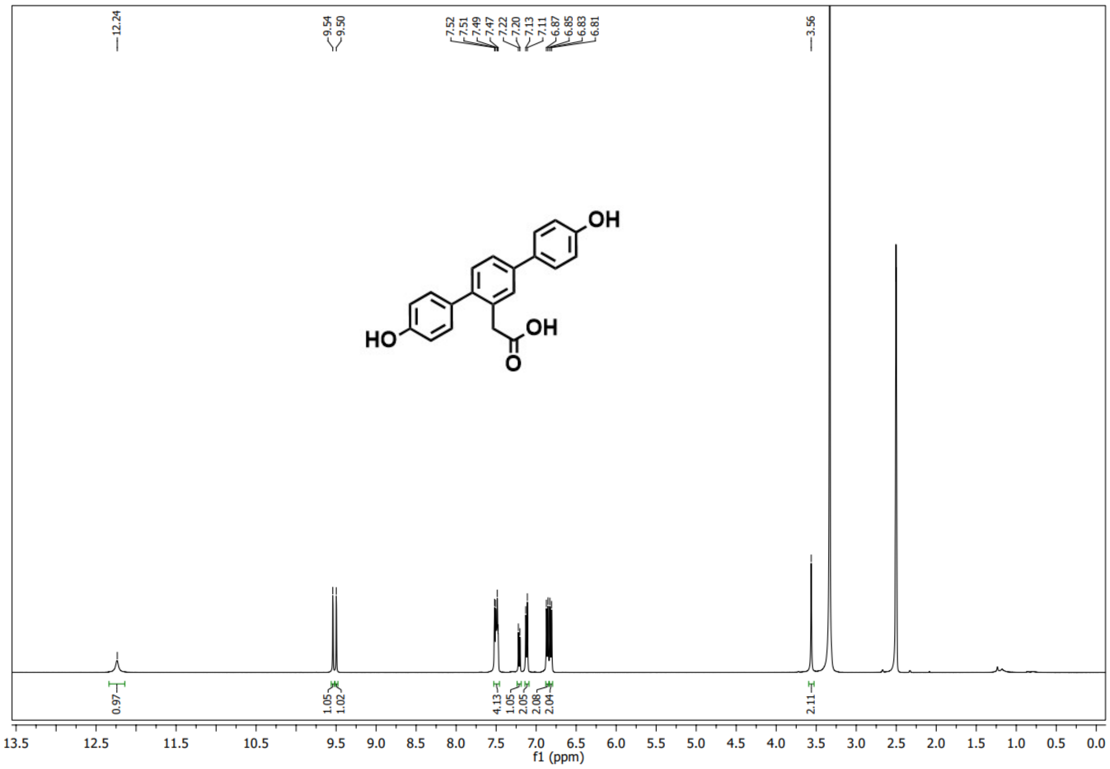

$^{13}$CNMR of **57-112** in DMSO-$d_6$

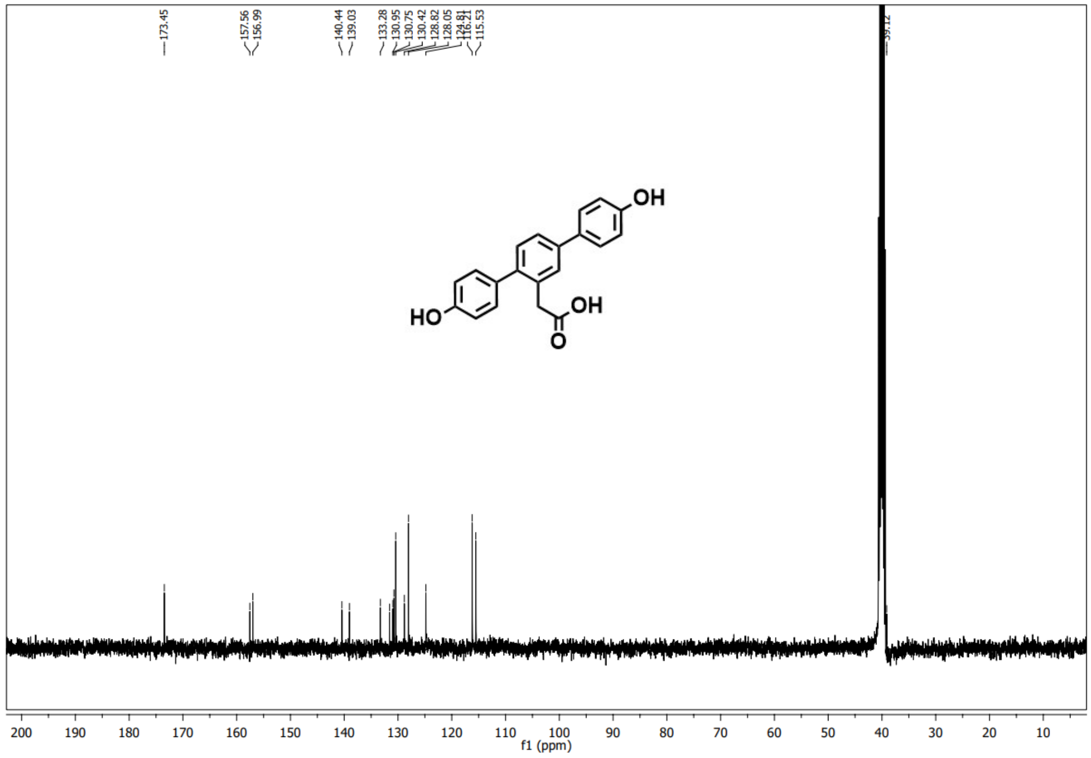

HPLC analysis of **57-112**

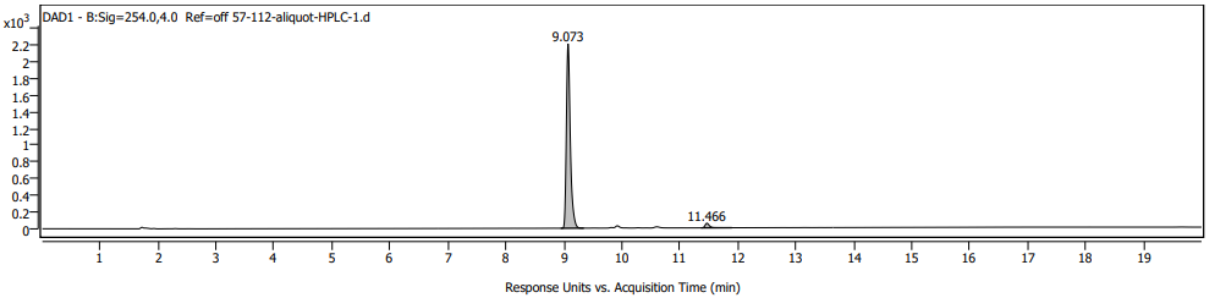


$^{1}$HNMR of **57-125** in DMSO-$d_6$

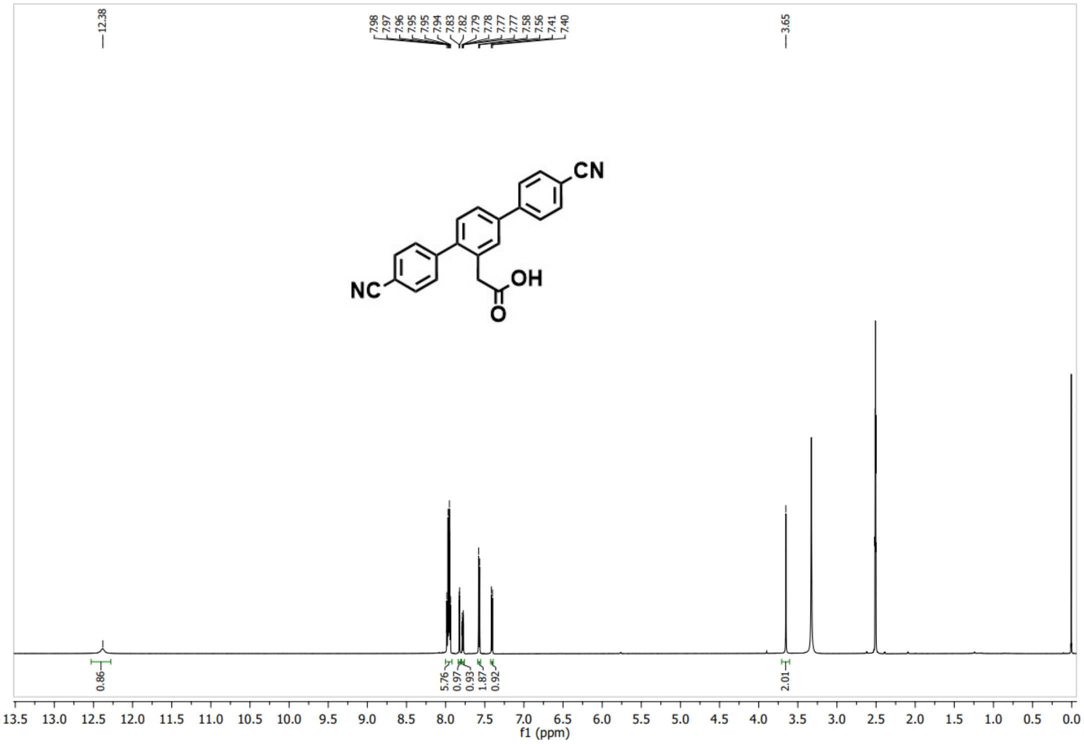


$^{13}$CNMR of **57-125** in DMSO-$d_6$

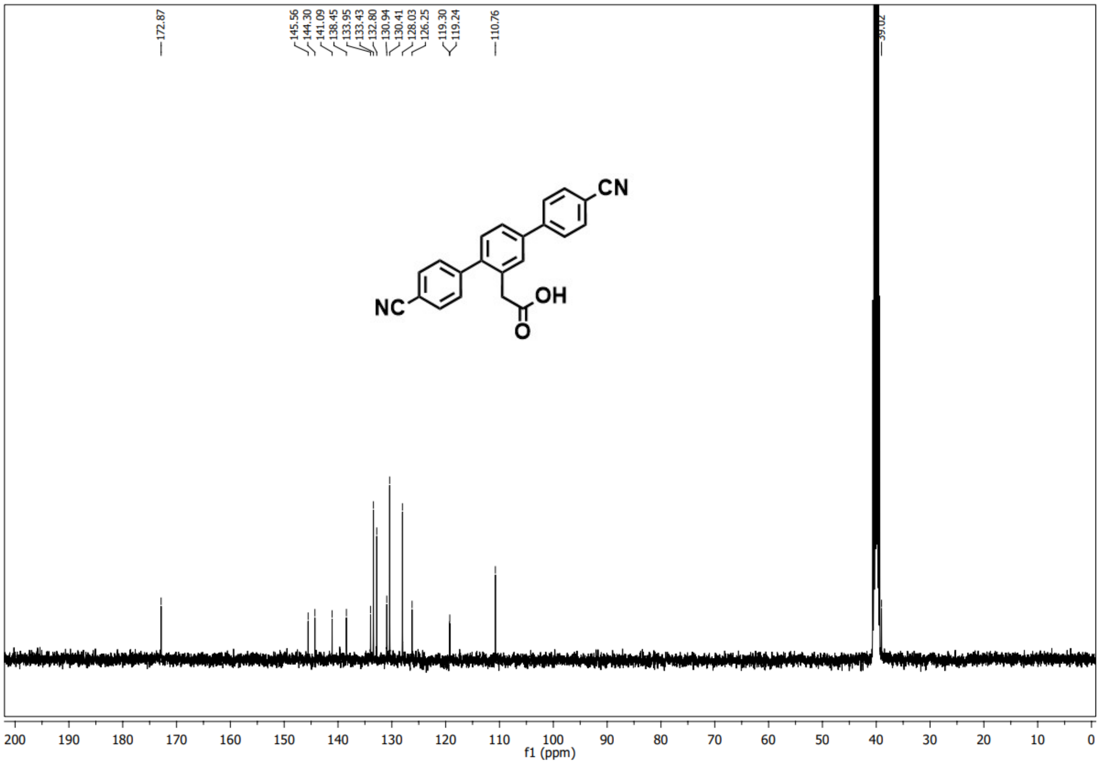


HPLC analysis of **57-125**

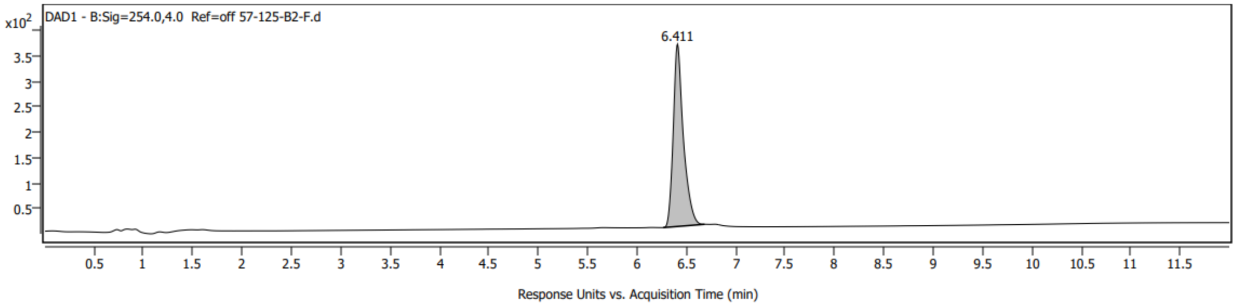


$^1$HNMR of **57-139** in $CDCl_3$

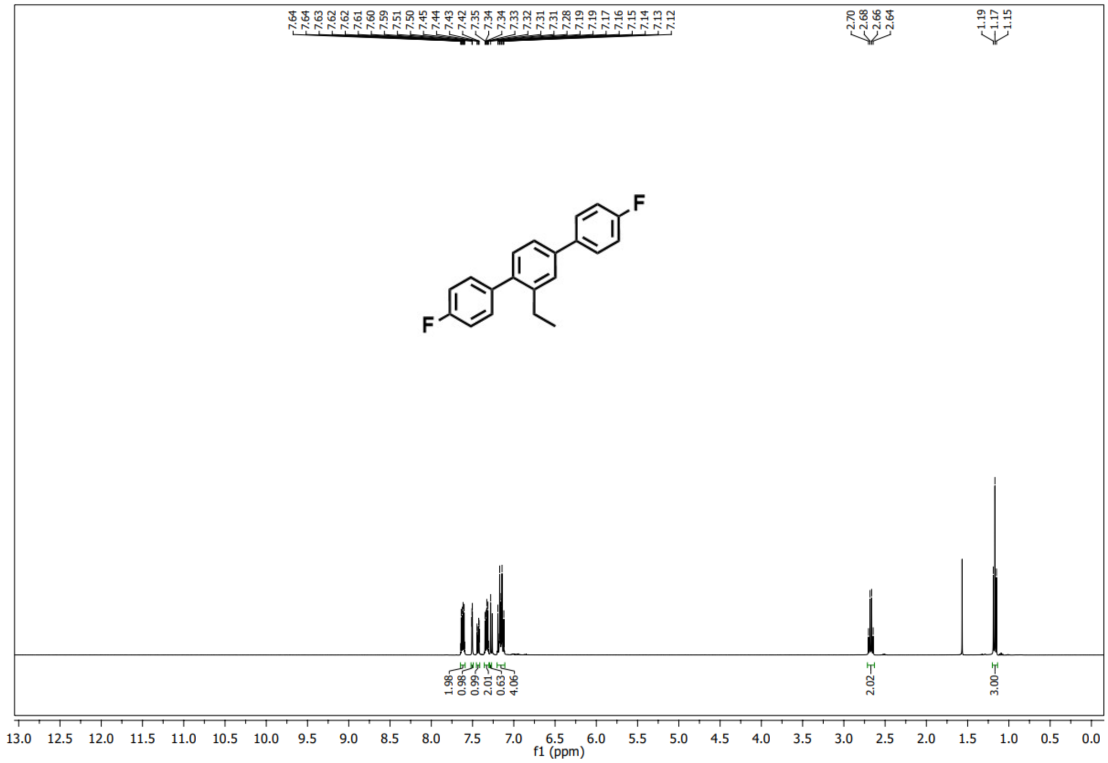


$^{13}$CNMR of **57-139** in $CDCl_3$

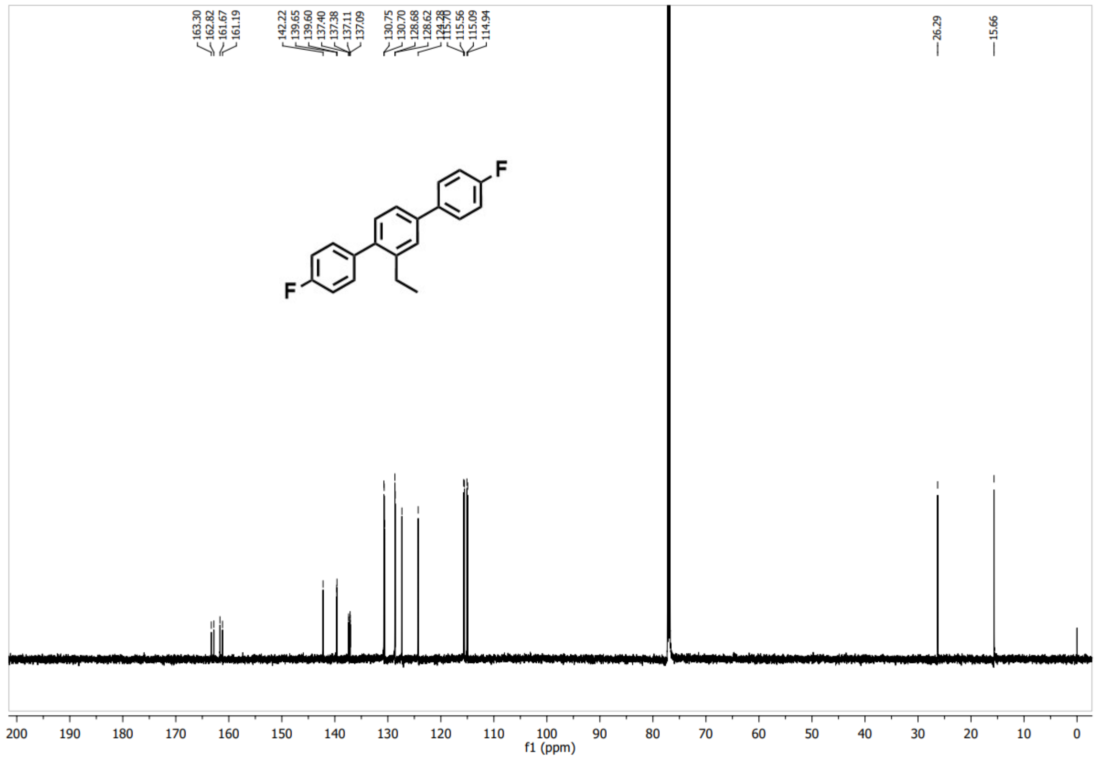


$^{19}$FNMR of **57-139** in $CDCl_3$

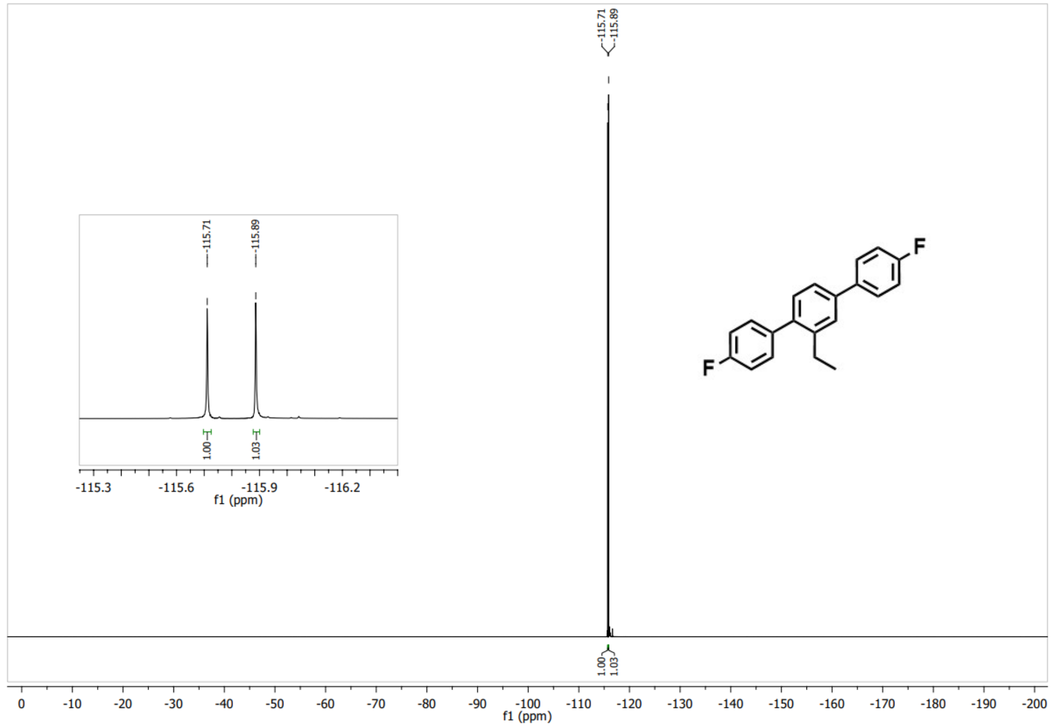


HPLC analysis of **57-139**

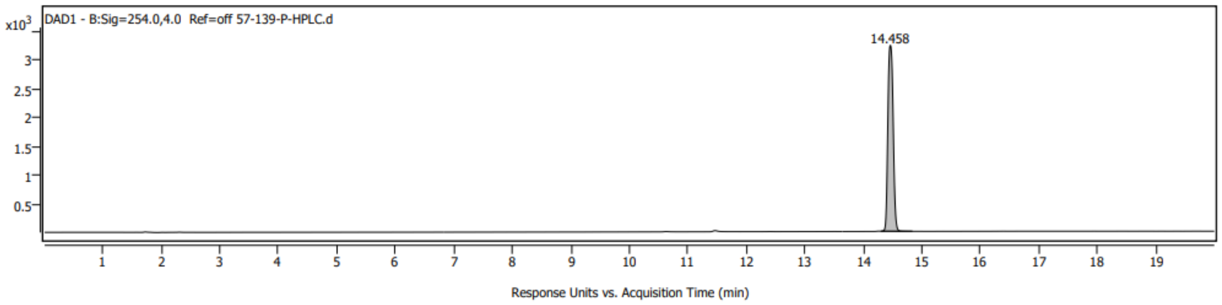
